# Simulation Insights on the Compound Action Potential in Multifascicular Nerves

**DOI:** 10.1101/2024.10.16.618681

**Authors:** Joseph Tharayil, Ciro Zinno, Filippo Agnesi, Bryn Lloyd, Silvia Farcito, Antonino M. Cassara, Niels Kuster, Michael Reimann, Silvestro Micera, Esra Neufeld

## Abstract

**Objective:** Develop an efficient method for simulating evoked compound action potential (eCAP) signals from complex nerves to help optimize and interpret eCAP recordings; validate it through comparison with measured vagus nerve eCAP recordings; elucidate the subtle interplay giving rise to specific eCAP signal shapes and magnitudes.

**Approach:** We developed an extended reciprocity theorem approach to model neuron signals in heterogeneous environments, and use it to study analytically the single fibre action potential. We then established a semi-analytic model that also uses hybrid electromagnetic-electrophysiological simulations to model eCAP signals from complex nerves populated with heterogeneous fiber populations of fibers. A cuff electrode was used to measure activity induced by vagus nerve stimulation in *in vivo* porcine experiments; these measurements were compared with signals produced by the model.

**Main Results:** The semi-analytic model produces signals that approximate the shape and amplitude of *in vivo* measurements. Partially activated fascicles contribute substantially to the signal, as eCAP contributions from smoothly varying fiber calibers in fully activated ones partially cancel. As a result, eCAP magnitude does not depend monotonically on the stimulation current and recruitment level. Because the eCAP is sensitive to the degree of activation in individual fascicles, and to the location of the recording electrodes with respect to individual fascicles, the contributions of different fascicles to the recorded eCAP signals vary significantly with changes in the shape and placement of the stimulus and the recording electrodes.

**Significance:** Our method can be used to rapidly assess new stimulation and recording setups involving complex nerves and neurovascular bundles, e.g., to maximize signal information content, for closed-loop control in bioelectronic medicine applications, and potentially to non-destructively reconstruct structural and functional nerve topologies through inverse problem solving.

## 1. Introduction

The compound action potential (CAP) is the electric signal that can be picked-up near a nerve and results from propagating axonal action potentials (APs) [1]. It can readily be recorded from a cuff electrode, making it valuable for neural code elucidation, neural sensing, and closed-loop control applications in the field of bioelectronic medicine (‘electroceuticals’,i.e., neural interfaces for regulating (organ-)physiological function, and neuroprosthetics). However, frequently only qualitative changes in, e.g., electrically evoked CAP (eCAP) magnitude are interpreted and the detailed eCAP shape and quantitative features are not leveraged. It is, for example, assumed that eCAP magnitude is monotonically related to the level of axonal recruitment [2].

Vagus nerve stimulation (VNS) is a promising bioelectronic medicine technique, which has already been approved for epilepsy and depression, and is under investigation for many other conditions [3]. Of particular interest are potential applications of VNS to cardiac disorders, including cardiac arrest, heart failure, and arrythmias [4].

The HORIZON 2020-funded NeuHeart project was launched with the aim of restoring autonomic control over cardiac function after heart transplantation with the help of a regenerative neural interface on the cardiac branch of the vagus nerve (VN), which both facilitates re-connection of the heart to the autonomic nervous system and permits to stimulate and regulate corresponding neural input [5]. NeuHeart also includes the development of a series of sensors for closed-loop stimulation control [6]. Computational modeling of implantable neural interfaces plays a central role in designing them, optimizing stimulation parameters, establishing model-based control strategies (where it is combined with an advanced cardiovascular regulation model [7, 8, 9]), and interpreting neural signals. eCAP signals are of interest to NeuHeart because 1) they can be used to validate model predictions on fiber recruitment by the implanted stimulation electrodes and 2) they could provide valuable additional feedback for closed-loop control.

The goals of this study were to:

- develop the *extended reciprocity theorem* approach as analytical basis for the accurate and efficient modeling of neural signals resulting from spatially extended neural activities in models of complex nerves with realistic neural interface designs;
- build a realistic semi-analytic model of the cardiac VN to support the NeuHeart project;
- compare predictions from that model against data from *in vivo* porcine experiments; and
- leverage the developed methodologies to improve understanding of the eCAP signal.

Peña et al. recently published a semi-analytic model of the eCAP [10], which leverages a similar approach to the one we introduced in [11]. Here, we extended it to multi-fascicular nerves with realistic, heterogeneous afferent and efferent, myelinated and unmyelinated fiber populations and spatially varying diameter probability distributions. We show that these factors critically influence the eCAP. This refined model is applicable to studying complex, multi-fascicular nerves and neurovascular bundles. The results were experimentally validated.

## 2. Methods

### 2.1. Mathematical model

We derived two related mathematical models: 1) an *analytical* model, described in Section 2.1.2 and 2.1.3, which predicts the single-fiber AP (SFAP) of an activated fiber as a function of the fiber properties and the recording electrode potential (both approximated by closed-form functions), and 2) a *semi-analytic* model (Sec. 2.1.2), which predicts the eCAP based on numerically computed characteristics of fiber models, an electromagnetic (EM) simulation of current applied to the recording electrodes, histology derived distributions of different fiber populations, and fiber recruitment curves obtained through coupled EM-electrophysiologial simulations. The image-based nerve model construction was performed in Sim4Life v7.2 (ZMT Zurich MedTech AG, Switzerland) The EM simulations, fiber functionalization, electrophysiology simulations, and extraction of quantities-of-interest were performed in Sim4Life v8.0.

#### 2.1.1. Extended Reciprocity Theorem

The well known reciprocity theorem states that the signal (voltage, *V* ) recorded between two electrodes resulting from a dipolar current source 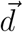 is related to the electric (E-)field generated at the dipole location 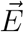 by (virtually) applying a current *J* to the same electrodes:

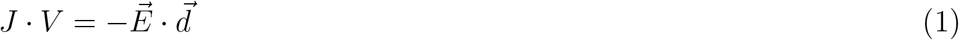

Consequently, simulation-based predictions of the signals generated by a series of time-varying dipole sources 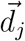 at locations *x*_*j*_ only require a single EM simulation of current applied to the sensing electrodes to evaluate 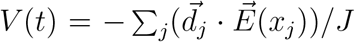 . For a neural fiber modeled as a line with a transmembrane current density *i*(*l*) along its length, it can be shown (c.f. Appendix A.1) that

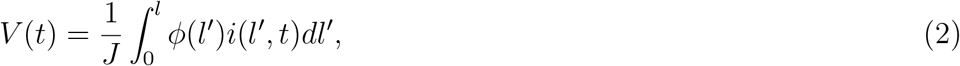

where *ϕ* is the electric potential along the fiber. We call Φ(*l*) = *ϕ*(*l*)*/J* the “sensitivity function” and refer to Equation 2 as the *generalized reciprocity theorem*. As a neuron with complex morphology can be decomposed into its line-shaped branches, that formula also readily applies, e.g., to pyramidal neurons with their dendritic arbors.

#### 2.1.2. Analytical SFAP Model

The trans-membrane current for a particular fiber type (index _*τ*_ ) can be obtained as 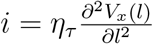 , where 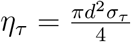 , *σ* is the axial conductance of an axon, *V*_*x*_ is the transmembrane voltage (distinguished by the subscript *x* from the temporal transmembrane potential *V*_*t*_(*t*)), and *d* is its diameter ([12]), see Sec. 4.2.2 regarding myelinated fibers).

From the extended reciprocity theorem (Eqn.(2)), it follows that the SFAP signal, *S*(*t*) is the convolution in space of the transmembrane current with the extracellular potential along the neuron. With the change of variables *x* = *t*^*′*^ · *v* (where *v* is the conduction velocity), we can express this convolution in the time domain as:

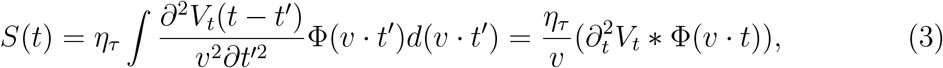

where ‘*’ signifies convolution in time. For a full derivation, see Appendix A.2. Taking the time Fourier transform ℱ _*t*_( ), we obtain

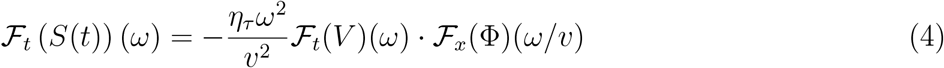

and through inverse Fourier transform 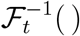 we recover the SFAP.

For myelinated fibers, conduction velocity is proportional to diameter [13], while for unmyelinated fibers, conduction velocity is proportional to the square root of diameter [14]. For the myelinated fiber model used in this study, the proportionality constant *a* = *d/v* was determined to be ∼ 4.3 · 10^6^*s*^−1^ (determined for a 20 *µ*m fiber; see Sec. 3.1, Sec. Appendix B and Fig. B1). For the unmyelinated fiber model, the proportionality constant 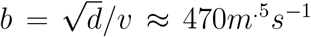 (based on 0.8*µ*m fiber). For convenience we define *η*_*τ*,*v*_(*d*) = *η*_*τ*_ */v*.

#### 2.1.3. Gaussian Approximation

To facilitate the analytical investigation of the eCAP contribution dependence on various variables, we approximate the AP shape as 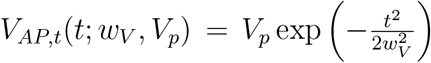 and the potential near a local monopolar electrode as 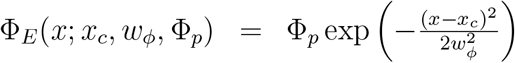, where *w*_*V*_ is the temporal AP width, *w*_*ϕ*_ the spatial electrode potential width, and *V*_*p*_ and Φ_*p*_ are the amplitudes. A bipolar electrode can be approximated as the combination of two monopolar contacts: Φ_*E*_(*x*; *x*_*c*,1_, *w*_*ϕ*,1_, Φ_*p*,1_) − Φ_*E*_(*x*; *x*_*c*,2_, *w*_*ϕ*,2_, Φ_*p*,2_).

Using Equation 4 we obtain:

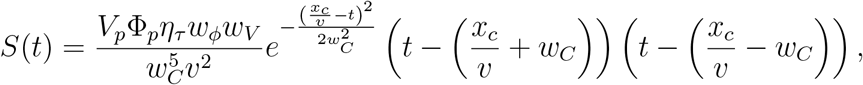

where 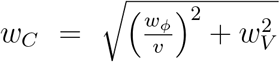 is the temporal Gaussian width after convolution and evidently the characteristic time-scale of the recorded signal. The SFAP shape corresponds to the typical peak between two side-lobes of opposite polarity – a result of the second derivative in Equation 3 – that peaks at *x*_*c*_*/v* – the time it takes to reach the recording electrode. The peak magnitude is:

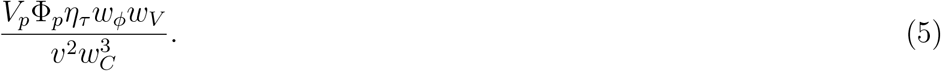

#### 2.1.4. Semi-Analytical eCAP Model

In a real nerve, a large number of fibers are simultaneously active. To determine the resulting eCAP signal, their SFAP contributions must be combined. Because of the linearity of Maxwell’s equations, simple additive combination can be used. To avoid simulating SFAPs from each fiber in the nerve, we construct the eCAP based on a representative SFAP per fiber type (index _*τ*_ ): myelinated afferents and efferents, and unmyelinated afferents and efferents. Electrophysiologically, afferents and efferents are identical.

For each fascicle (index _*i*_), we consider 2000 fibers of each fiber type, with uniformly spaced diameters from 0.1 to 15 *µ*m. Each SFAP is scaled by the fiber diameter density *n*_*i*,*τ*_ (*d*) = *N*_*i*,*τ*_ ·*p*_*τ*_ (*d*) (*N*_*i*,*τ*_ : number of fibers of the given type, *p*_*τ*_ (*d*): diameter probability distribution) and the degree of fascicle recruitment *R*_*i*,*τ*_ (*d*) (Fig. 4.a-b). Note that *R*_*i*,*τ*_ (*d*) implicitly depends on the stimulation magnitude and pulse shape.

The fiber number *N*_*i*,*τ*_ is a random variable drawn from distributions (obtained by [15] based on *in vivo* data) which are dependent on the location of the fascicle within the nerve (myelinated afferents and efferents are localized on opposite sides of the nerve, and unmyelinated efferents are colocalized with myelinated afferents), and scaled by the area of the fascicle. For details on the calculation of *R*_*i*,*τ*_ , see Section 2.3.3.

In multifascicular nerves, the different fascicles are separated by semi-insulating perineurium, such that within fascicle *i* the *ϕ*_*i*_(*l*) along fiber trajectories is similar for all fibers, but the eCAP contributions must be computed using different *ϕ*_*i*_(*l*) for the different fascicles (Fig. 4.e).

Thus, the eCAP is given by

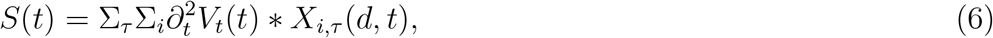

where the fascicle “exposure function” *X*_*i*,*τ*_ (*d, t*) is defined as 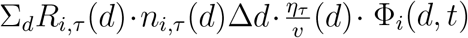 and Δ*d* is the width of the discretized fiber diameter bins (see Appendix H).

### 2.2. Experimental Data Acquisition

Experimental data was acquired from minipigs. The surgical protocol is described in Appendix C. The NeuHeart Vagal Regenerative Autonomic Interface (VRAI) electrode [16], which is a transversal, intrafascicular, multichannel electrode (TIME) featuring eight contacts on each side was inserted in the cardiac branch of the VN for stimulation purposes, and a cuff electrode with four ring contacts was placed at a distance of either 1 or 6 cm from the stimulation electrode to record eCAP signals. Varying stimulation intensities (100 to 500 *µA*), active contacts, and pulse repetition rates (1 to 50 Hz) were applied. The experimental presented in this paper have all contacts active, and use either a 50 Hz (Fig. 3.a) or 30 Hz (Fig. 3c). pulse repetition rate. Note that in our simulations, we do not model the effect of pulse repetition. Stimulus-triggered averaging was used to reduce noise. Additional details about the experimental methods are provided in Appendix C.

### 2.3. Realistic, Multifascicular Nerve Model

To demonstrate the application of the extended reciprocity theorem and of the semi-analytical modeling in a realistic setup, we modeled the experimental setup from Section 2.2.

#### 2.3.1. Anatomical Geometry

A 2D cross section of the pig VN was segmented from a histology slice, the tissue contours were extruded to produce a 2.5D model, and a model of the stimulating VRAI and the recording cuff electrode was integrated (see Fig. 1).

**Figure 1.**
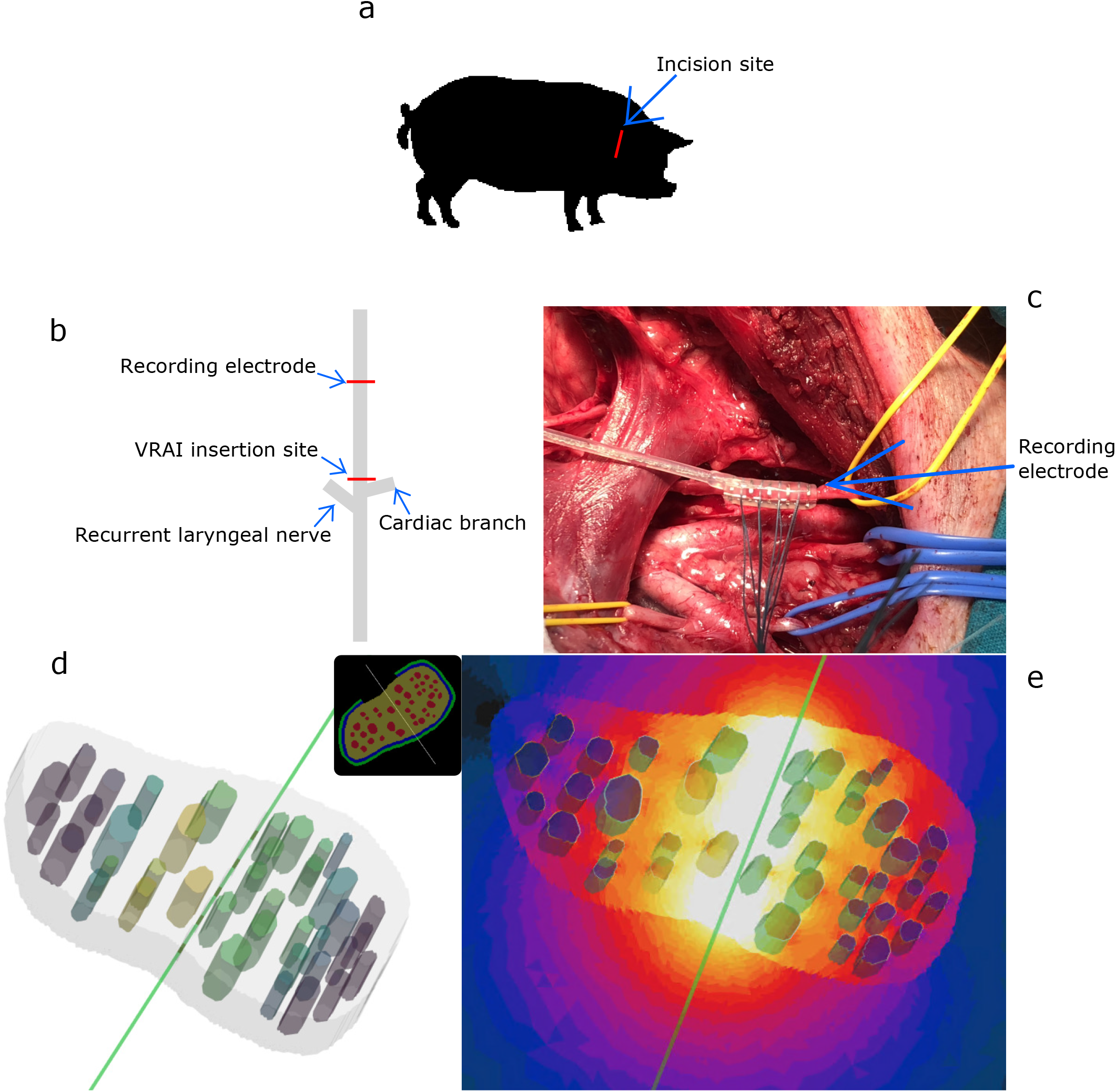
a: Incision site on the minipig. b: Schematic of the VN. c: Photo taken during surgery, showing the recording electrode on the VN. d: VN model with NeuHeart VRAI stimulation electrode (inset: nerve cross section surrounded by the cuff recording electrode). Note that the histology-based cross section topology likely differs from that of the experimentally stimulated nerve (different minipigs). The placement of the VRAI is one of two alternatives studied in this paper. Except where otherwise noted, modeled eCAPs are based on the insertion trajectory from Fig. 5.c). e: E-field from the electrode. Fascicles are clearly visible due to the insulating effects of the perineurium.

#### 2.3.2. EM Modeling

The complete model was discretized, and the Sim4Life Electro-Ohmic Quasi-Static (EQS) solver was used to calculate potentials and E-fields resulting from an electrical stimulus applied at the VRAI electrodes and from a virtual current applied at the cuff electrodes (c.f. Appendix D for details).

#### 2.3.3. Neuronal Dynamics Simulations

The temporal profile of the AP was obtained from a multicompartmental simulation in Sim4Life of a rat myelinated sensory-motor neuron [17, 18] and a Sundt unmyelinated neuron [19], for myelinated and unmyelinated fibers, respectively.

The recruitment *R*_*i*_ for a given stimulus waveform is calculated as follows: In our finite element model of the VN, we instantiate a population of myelinated fibers with uniform diameter *d*_0_ = 4 *µ*m in each fascicle and a unmyelinated population with *d*_0_ = 0.8 *µ*m. For each fascicle, a recruitment curve 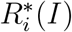 was generated by titrating simulated VRAI field induced nerve activation. Thanks to the quasi-linear diameter-dependence of fiber stimulation thresholds, 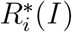 can be recast as diameter-dependent *R*_*i*_(*d*) for a given stimulus of intensity *I* according to 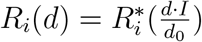 .

## 3. Results

### 3.1. SFAP Shape in Simple Analytical Model

The propagation velocity *v* is strongly fiber diameter- and type-dependent. Two limit cases of the Gaussian approximation model are of interest: When *v* ≫ *w*_*ϕ*_*/w*_*V*_ (i.e., the AP extent is much larger than the region to which the sensing electrode is sensible), the temporal width *w*_*C*_ ≈ *w*_*V*_ and the amplitude becomes . 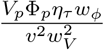 When *v* ≪ *w*_*ϕ*_*/w*_*V*_ , the temporal width *w*_*C*_ */v* and the amplitude becomes 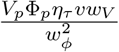 (Table 1). For the recording electrodes used in this paper, *w*_*ϕ*_ ∼ 1 mm (Fig. 8.a). For the myelinated fiber model used in this study, *v* ∼ 4.3 <u>·</u> 10^6^*d* and *w*_*V*_ ∼ 5 · 10^−5^s (Fig. B1.a,c). For the unmyelinated fiber model, 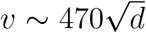 and *w*_*V*_ ∼ 2 · 10 s^−4^ (Fig. B1.b,d). Thus, for the unmyelinated fiber model, realistic fiber diameters are in the *v* ≪ *w*_*ϕ*_*/w*_*V*_ regime the theoretical diameter *d*^*^ at which *v* = *w*_*ϕ*_*/w*_*V*_ is *>*100 *µ*m. For myelinated fibers, however, *d*^*^ ≈4.7 *µ*m, which is comparable in scale to the diameters of the relevant fiber population, such that their eCAP contribution show a more complex *d* dependence in terms of magnitude, but also shape (see Sec. 4.1).

**Table 1.** Velocity as a function of diameter (first column) and diameter *d*^*^ at which , 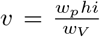 for *w*_*ϕ*_ = 1 mm, (second column) for myelinated and unmyelinated fibers. Maximum amplitude of SFAP for myelinated and unmyelinated fibers, in general (third column) and in the two limit cases (AP extent much shorter or longer than electrode potential; fourth and fifth columns). 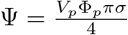.

| $v$ | $d^*$ | | general | $d \ll d^*$ | $d \gg d^*$ |
| --- | --- | --- | --- | --- | --- |
| | | | $w_C: \sqrt{\left(\frac{w_\phi}{v}\right)^2 + w_V^2}$ | $w_\phi/v$ | $w_V$ |
| myelin.: $4.3 \cdot 10^6 d$ | $4.7 \mu\text{m}$ | amplitude: | $\frac{\Psi w_\phi w_V}{a^2 w_C^3}$ | $\frac{\Psi a w_V}{w_\phi^2} d^3$ | $\frac{\Psi w_\phi}{a^2 w_V^2}$ |
| unmyel.: $470\sqrt{d}$ | $110 \mu\text{m}$ | amplitude: | $\frac{\Psi d w_\phi w_V}{b^2 w_C^3}$ | $\frac{\Psi b w_V}{w_\phi^2} d^{5/2}$ | $\frac{\Psi w_\phi}{b^2 w_V^2} d$ |

As a result of the square-root dependence of the unmyelinated fiber conduction velocity on diameter, their SFAP contribution magnitude monotonously increases with diameter (see Table 1 and Fig. 2.a). For the linearly dependent myelinated fibers, however, the contribution weight saturates once *d* ≫ *d*^*^ ≈ 4.7*µm* (Table 1, Fig. 2.b). This model-based analysis uses strong simplifications (e.g., shape of the AP) and the SFAP magnitude dependence on fiber diameter is subtle, such that for real-world fibers, it could even be maximal for an intermediate fiber caliber.

**Figure 2.**
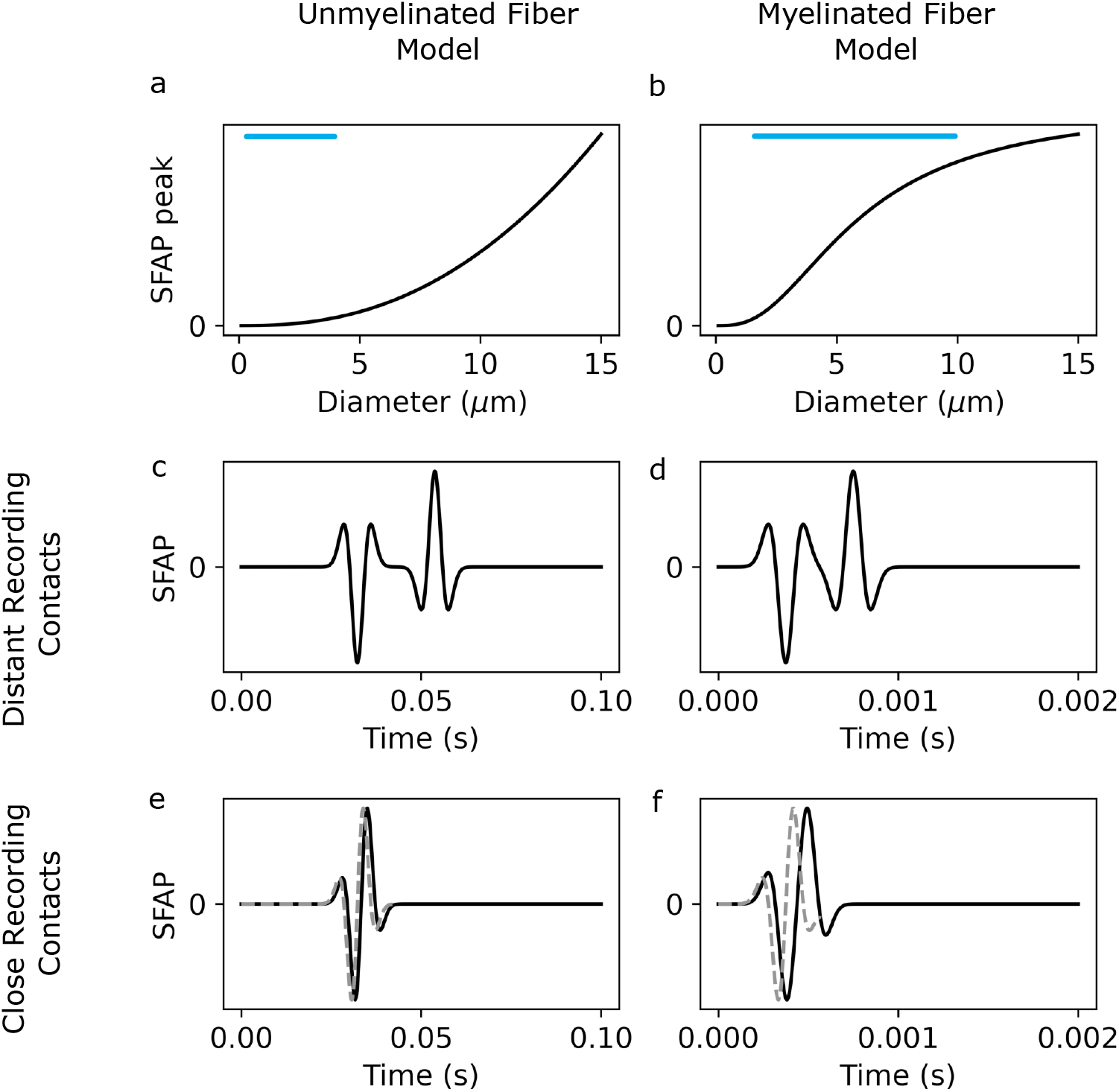
Properties and behavior of the SFAP in the analytical model. a,b: The SFAP amplitude increases with diameter for unmyelinated fibers but plateaus for the myelinated fibers. Blue bar indicates realistic fiber diameter range. c, d: Distinct trimodel waveforms are apparent for sufficiently separated contacts in the bipolar recordings of an unmyelinated fiber (c; diameter: 1 *µ*m, spacing: 1 cm) and a myelinated fiber (d; diameter: 10 *µ*m, spacing: 1.5 cm). e,f: With decreasing contact separation, the two waveforms merge and a single tetramodal one results (e: unmyelinated fiber, spacing 0.1 cm spacing; f: myelinated fiber 0.5 cm spacing). It resembles the derivative of the original waveform (dashed).

In the case of the bipolar recording, when the two electrodes are sufficiently separated they produce an SFAP that simply reflects the distinct arrival of the AP at the two electrodes and their opposite polarities (Fig. 2.c-d). However, once 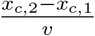 is in the same order of magnitude as *w*_*C*_, the two shapes start to overlap, which can result in fewer lobes. For unmyelinated fibers in the range of 0.1 to 4 *µ*m, the velocity *b* ·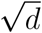 ranges from 0.15 to 0.93 m/s, which in combination with *w*_*V*_ results in a spatial AP extent in the order of 1 mm (Fig. 2.e). For myelinated fibers in the range of 1 to 10 *µ*m, the velocity *a* · *d* ranges from 4.4 to 44 m/s, which in combination with *w*_*V*_ results in the same spatial AP extent in the order of 1 mm (Fig. 2.f). This implies that an electrode separation of less than 1 mm will result in the merging of the SFAP waveforms. Because of the bimodal shape of *ϕ* in bipolar recordings, convolution with *ϕ* is akin to a finite difference derivative and produces a signal that resembles the first derivative of a monopolar recording (see Fig. 2 and Appendix I).

### 3.2. Semi-Analytic Model Replicates In Vivo eCAP

The semi-analytic model, with 6 cm stimulus-recording separation, is able to replicate key features of the *in vivo* eCAP. The overall shape of the *in vivo* eCAP (Fig. 3.a) – with an initial positive deflection, a negative deflection of similar width and magnitude, and a longer, lower-amplitude positive deflection – is reproduced by the model (Fig. 3.b). However, the second negative deflection at 5 ms is not reproduced. While the model replicates the saturation of the eCAP signal with increasing current, it does not replicate the difference in amplitude between the eCAP evoked at *I* =100 *µ*A and eCAPs evoked at greater amplitudes. The magnitude of the model eCAP is smaller than the *in vivo* signal by approximately a factor 2.

**Figure 3.**
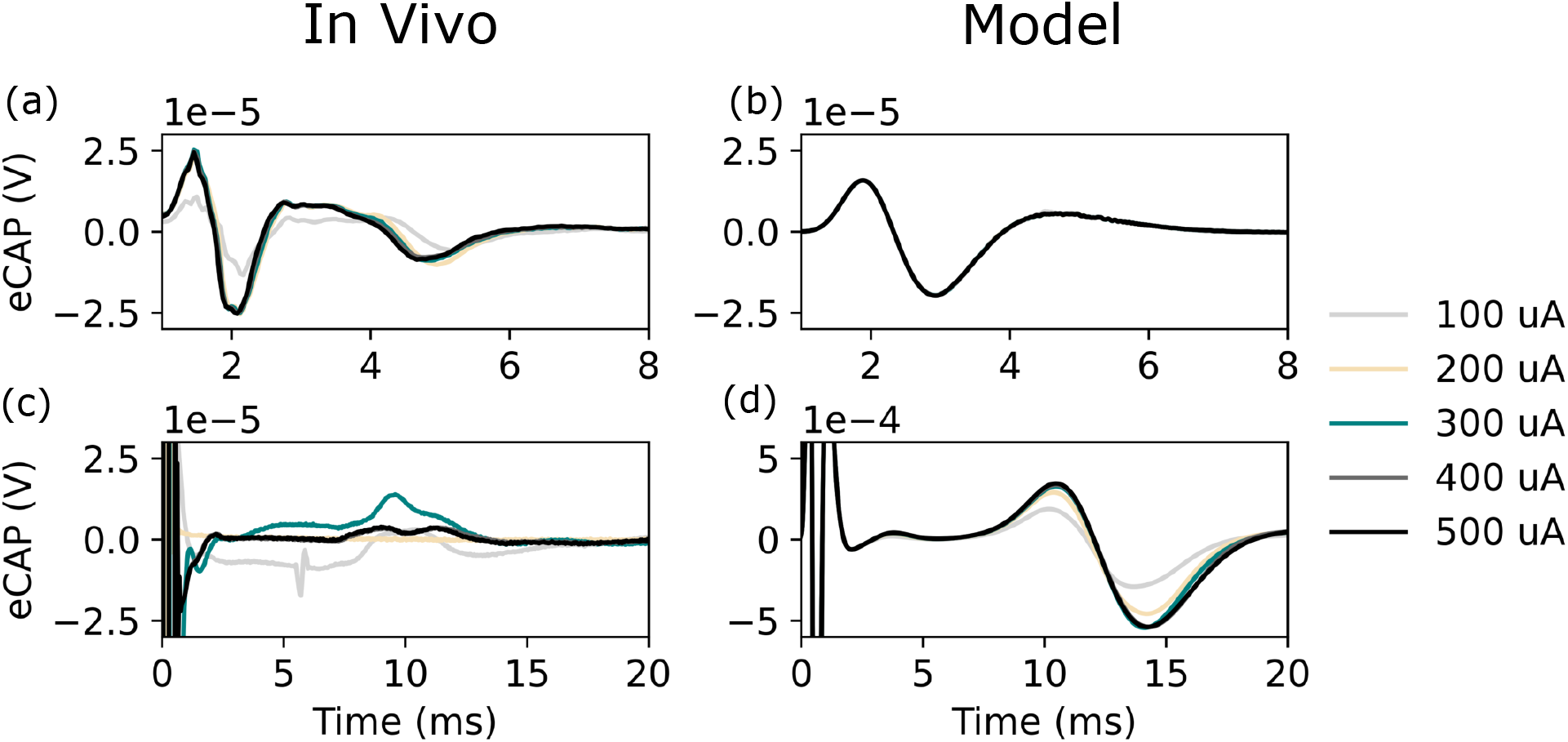
The semi-analytic model replicates the *in vivo* eCAP. Note that reported stimulus amplitudes refer to per-contact amplitude for the *in vivo* experiments, but to total amplitude summed over contacts for the model. a: *In vivo* eCAP recorded 6 cm from stimulus electrode. Signal begins at 1 ms to avoid stimulus artifact. b: Semi analytic model eCAP recorded 6 cm from stimulus electrode. Signal begins at 1ms for consistency with panel a. c: *In vivo* eCAP recorded 1 cm from stimulus electrode. d: Semi-analytic model eCAP recorded 1 cm from stimulus electrode.

With a distance of 1 cm between the stimulus and recording electrodes, the model (Fig. 3.d) replicates the timing of the secondary deflection observed *in vivo* (Fig. 3.c). Our model demonstrates that this secondary deflection is due to the activation of unmyelinated fibers. However, the magnitude of the deflection is not matched (see Discussion).

### 3.3. Fascicular Contributions to eCAP Cluster With Degree of Activation

To explore the influence of fiber activation on the eCAP, we reduce the stimulus amplitude to 31.25 *µ*A. The shape and amplitude of the fascicular contribution to the eCAP depend non-trivially on the degree of activation. For strongly activate fascicles (Fig. 4.a, orange curve), the exposure function *X*_*i*_(*t*) (Fig. 4.e), and correspondingly, the eCAP (Fig. 4.h) produced by convolution with the transmembrane current (Fig. 4.g) are smooth. For fascicles with an intermediate degree of activation (Fig. 4.a, blue curve), the exposure function (Fig. 4.f) is non-smooth, resulting, for the simulated configuration, in a large-amplitude deflection in the eCAP at 3.5 ms (Fig. 4.h). This is because the absence of activated fibers below 3 *µ*m means that the contributions of fibers above this threshold are not canceled out (see Sec. 4.1).

**Figure 4.**
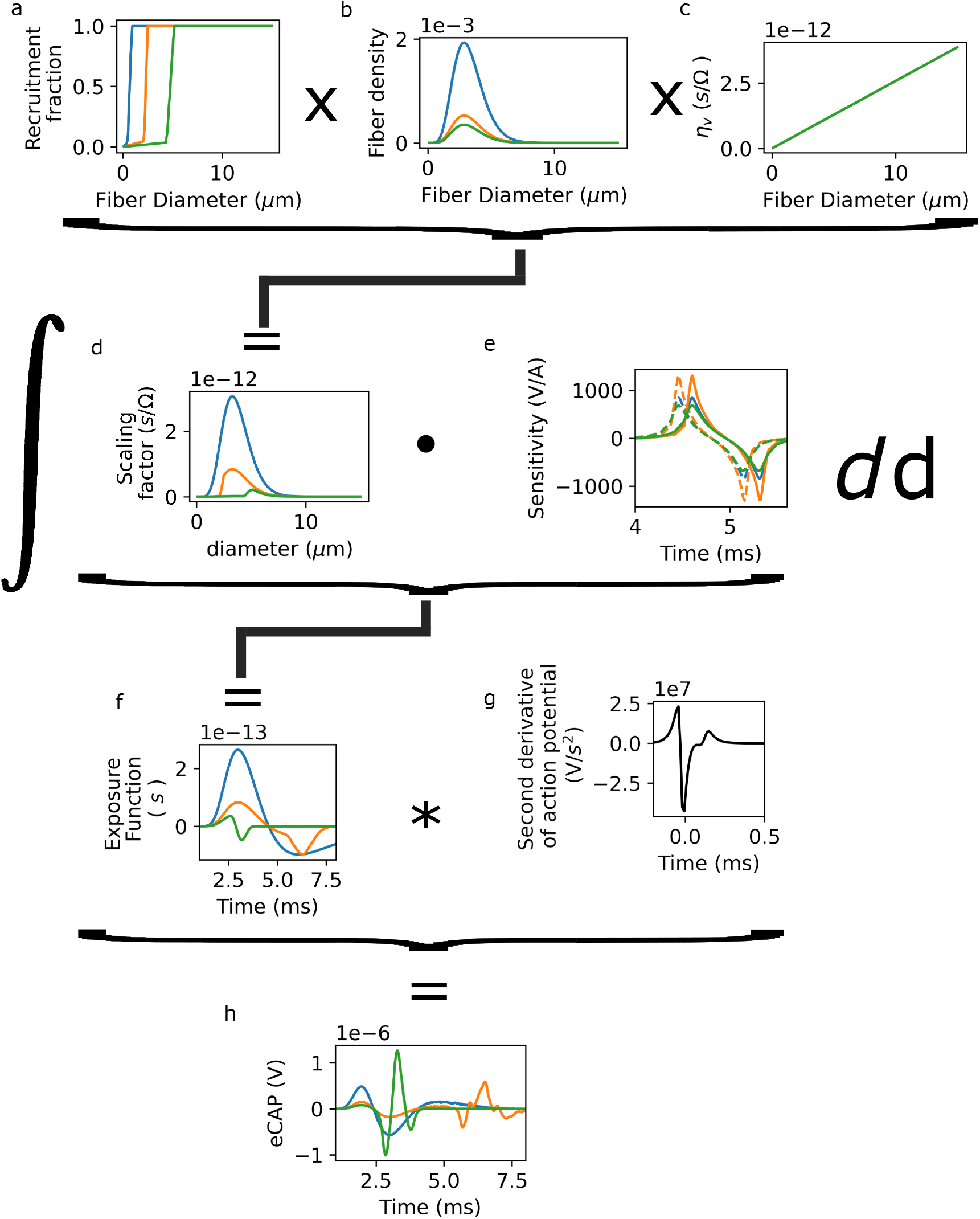
Calculation of the to the eCAP for myelinated afferent fibers from three representative fascicles (selected based on the clustering in Fig. 5; consistent color coding). a: Fraction of myelinated afferent fibers *R*_*i*_(*d*) recruited as a function of diameter. b: Myelinated afferent fiber diameter distribution *N*_*i*_ · *p*(*d*). c: Scaling 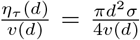 of transmembrane currents as a function of fiber diameter. d: Scaling factor 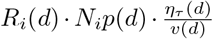Sensitivity function Φ_*i*_(*d, t*), plotted for diameters 3 *µ*m (solid lines) and 3.1*µ*m (dashed lines). f: Exposure function *X*_*i*_(*t*) for each fascicle. Sum of Φ(*d, t*) scaled by the diameter-dependent factor 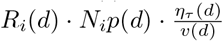 over all diameters. g: Second derivative of the AP; identical to the transmembrane current up to a scaling factor 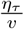 (panel c). h: eCAP contribution; convolution of f and g.

Therefore, the eCAP contributions from the individual fascicles are strongly correlated (Pearson product-moment correlation) for fascicles with similar degrees of activation (Fig. 5). The fascicles were clustered into three groups based on the correlations between their eCAP contributions, using k-means clustering. Fascicles in each cluster have similar spatial locations (Fig. 5.b), fractions of fibers activated (Fig. 5.c), recruitment curves (Fig. 5.d), and eCAP contributions (Fig. 5.e). We observed that fascicles in the center of the nerve, close to the stimulus electrode, belong to one cluster with high activation, steep recruitment curves, and smooth eCAP contributions.

**Figure 5.**
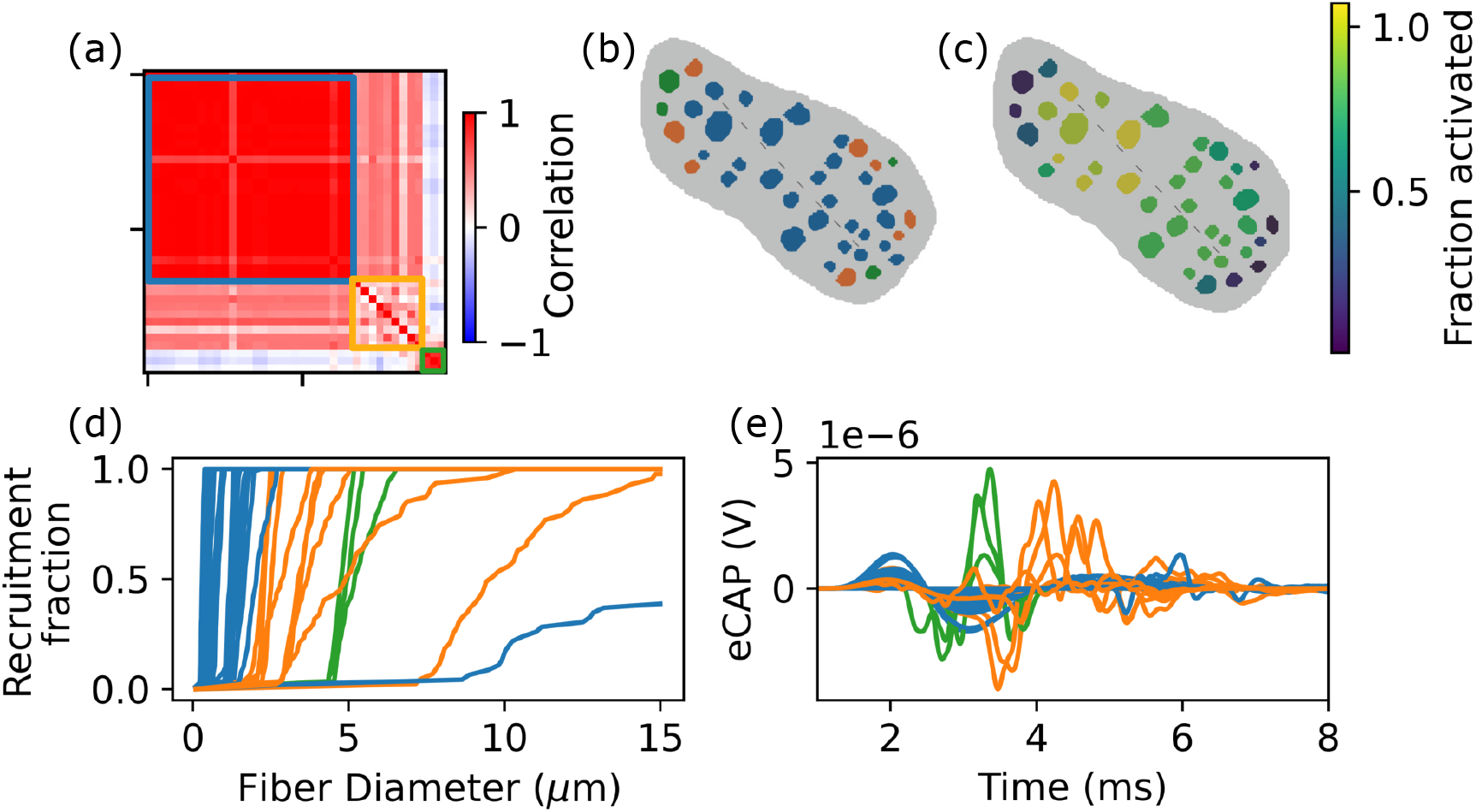
Fascicular recruitment dominates the eCAP contribution shape. a: Correlations between eCAP contributions between the different fascicles (stimulus amplitude: 31.25 *µ*A). Three clusters, identified using k-means, are outlined (cluster color coding consistent throughout Fig. 5 and 6). b: Cross section of the nerve, with fascicles colored by cluster. c: Fraction of myelinated fibers activated in each fascicle d: Cluster-colored recruitment curves for individual fascicles. e: Cluster-colored eCAP contributions for individual fascicles.

In contrast, fasiclces at the edges of the nerve, farther form the stimulus electrodes, have a lower activation fraction, and consequently, less smooth eCAP contributions.

The stimulus amplitude (*I*) dependent peak magnitude of the fascicular contributions to the eCAP follow a stereotyped pattern, rising to a maximum before decaying and plateauing once recruitment reaches 100% (Fig. 6.a,c). This can be explained by the non-monotonic relationship between fiber activation and eCAP ampltiude demonstrated in Figure 4. Consequently, the overall eCAP also does not always monotonically increase with fiber activation (Fig. 6.b). As expected, fascicles in the same cluster, as described above, have similar stimulus-eCAP amplitude relationships (Fig. 6.a).

**Figure 6.**
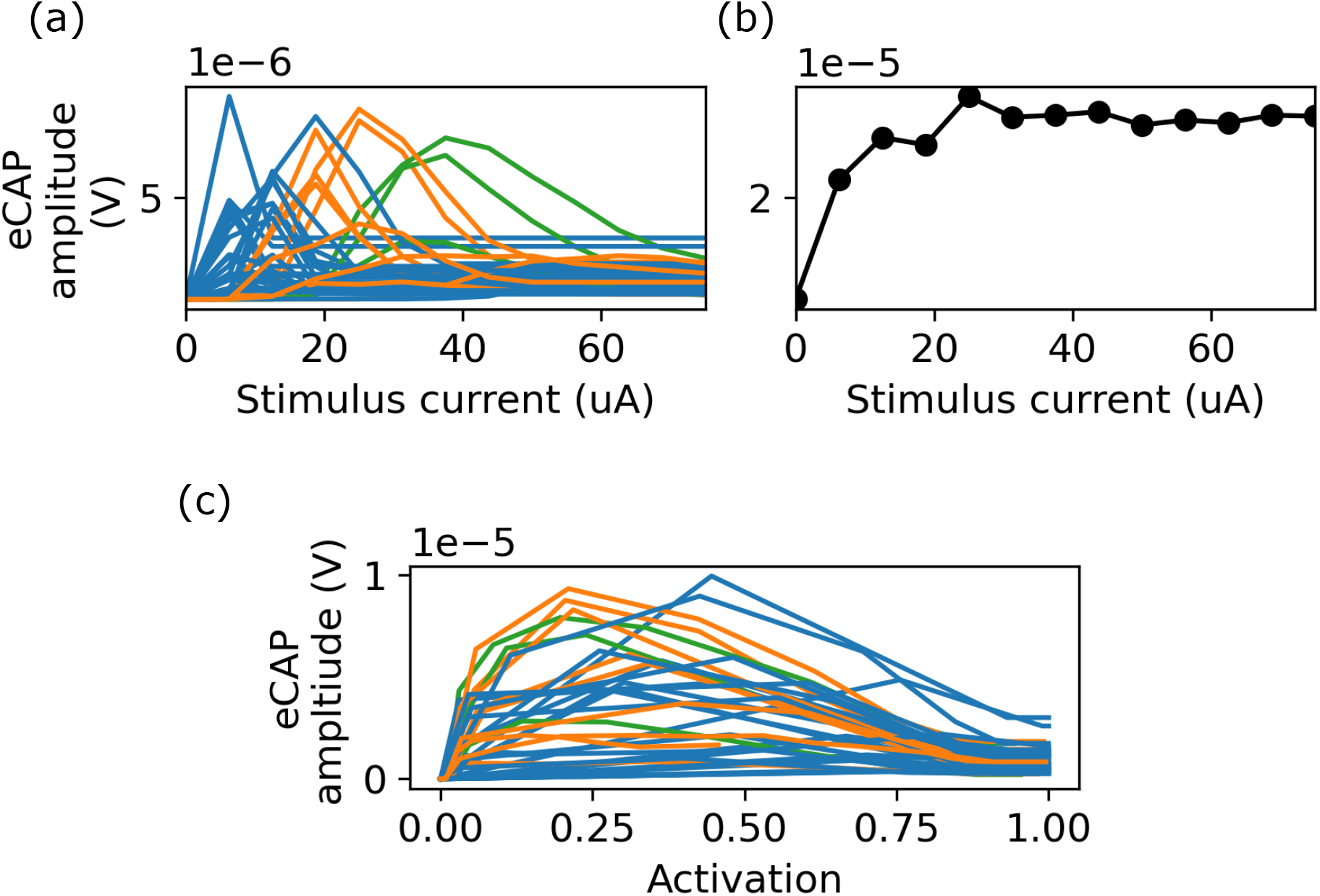
Non-monotonicity of fascicle recruitment dependent eCAP contribution magnitude. a: Peak amplitude of the contribution to the eCAP from each fascicle, as a function of current. Curves for each fascicle are colored according to the clustering from Fig. 5.b: Peak amplitude of the total eCAP, as a function of stimulus current. c: Peak amplitude of the fascicular eCAP contributions, as a function of their degree of activation.

### 3.4. Interactions Between Fascicular Structure and Stimulation Electrode Geometry Influence eCAP

The insertion angle of the stimulus electrode with respect to the nerve has a substantial impact on the eCAP (Fig. 7.a), as changes in the distance and orientation of the stimulus contacts relative to the different fascicles (Fig. 7.c,e) substantially change the degree of fiber activation therein (Fig. 7.b,d). Similarly, the orientation and extent of the recording electrode with respect to the nerve influences the eCAP (Fig. 8.a), because the distance from the electrode to each fascicle (Fig. 7.e, 7.g, 8.c) has a significant effect on each fascicle’s sensitivity function Φ_*i*_(*l*) (Fig. 7.d, 7.f, 8.b).

**Figure 7.**
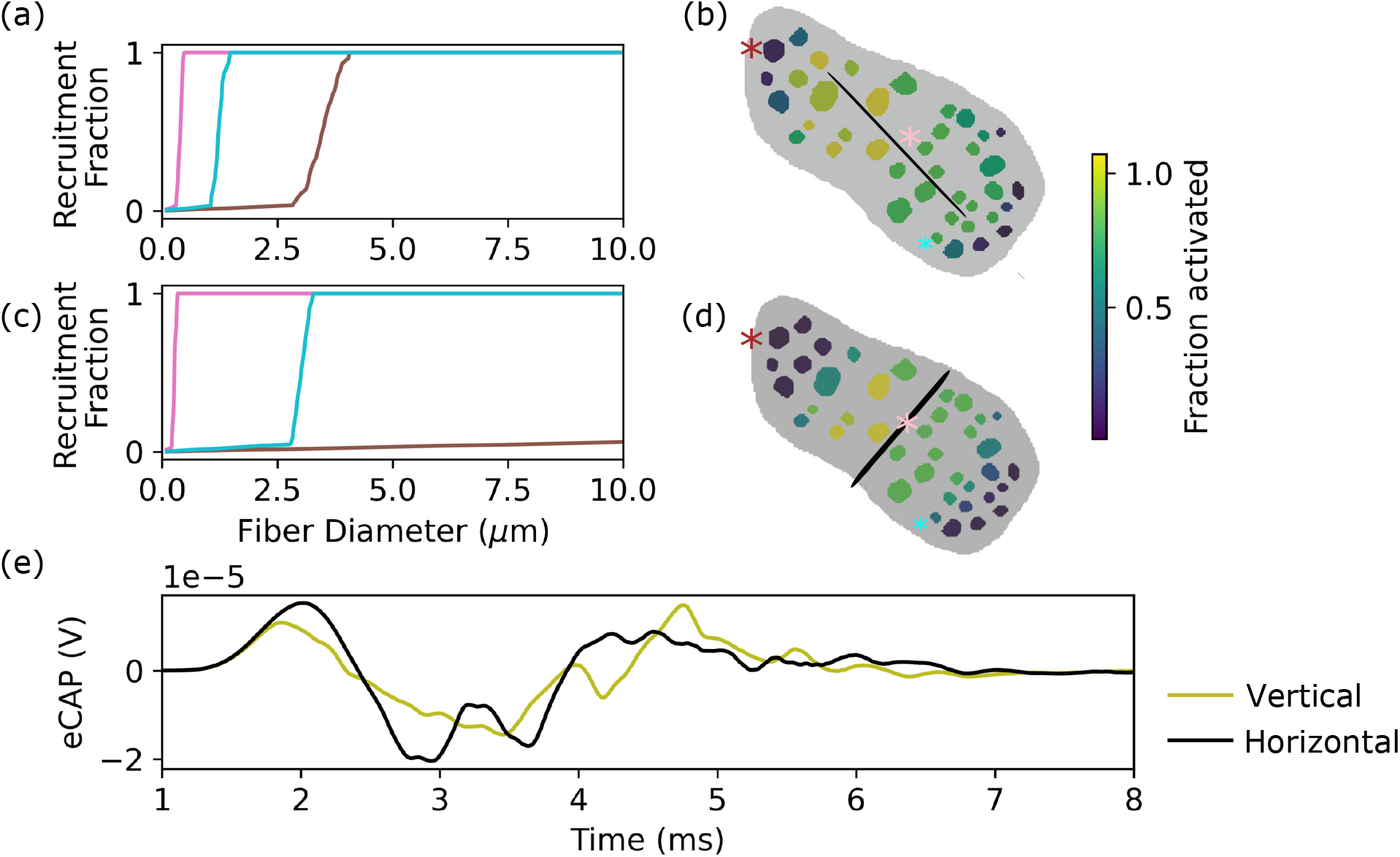
Stimulus electrode orientation influences eCAP due to differential activation of fascicles. a: Recruitment curves for selected fascicles, electrode inserted along the major axis of the nerve (stimulus amplitude: 31.25*µ*A) b: Fraction of myelinated fibers activated for each fascicle (electrode trajectory highlighted with black line for visualization purposes only; selected fascicles from panel a are marked with correspondingly colored asterisks). c-d: Same as a-b., for electrode insertion along the minor axis of the nerve. e: Simulated eCAPs.

**Figure 8.**
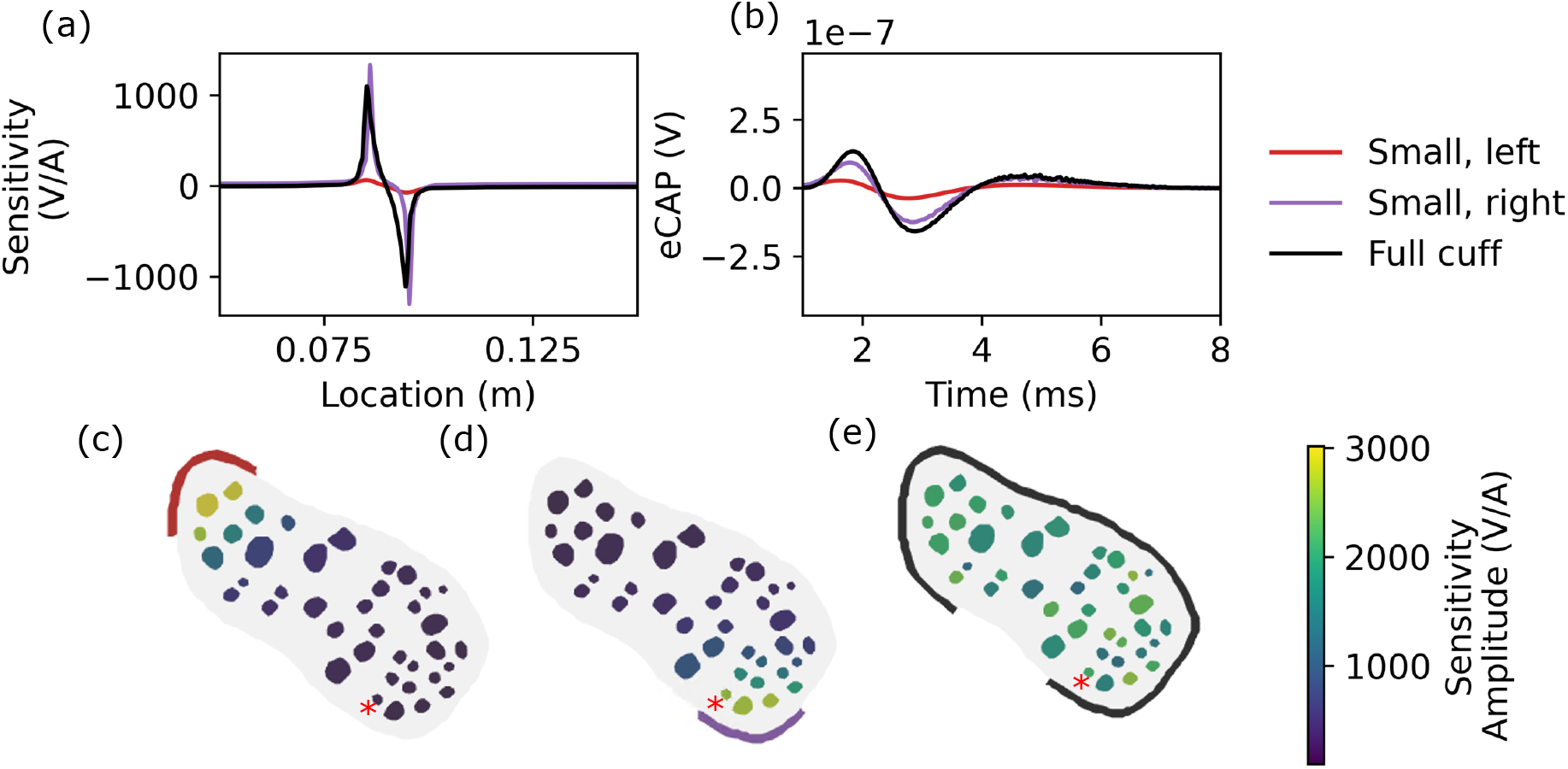
The size and orientation of the recording electrode influences the eCAP due to differing fascicular sensitivities Φ_*i*_. a: Sensitivity to activity in a selected fascicle (marked with red asterisk in panels c-e) for three different electrodes. b: Corresponding eCAP contributions. c-e: Peak-to-peak magnitude of sensitivity function, for each fascicle and recording electrode configuration.

## 4. Discussions

### 4.1. eCAP Cancellation and Partial Nerve Activation

Based on Eqn.(2), it can readily be shown that eCAP signal contributions subtly annihilate on the single fiber and fiber population level:

#### 4.1.1. Single Fiber

For propagating APs, Eqn.(2) can be restated as:

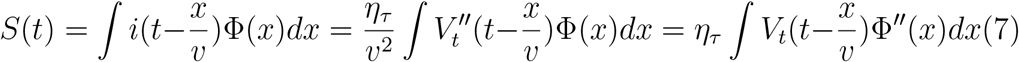

where *i*(*t*) and *V* (*t*) refer to the time dependent transmembrane current and voltage at reference location *x* = 0.

Different situations can be distinguished: If the AP’s spatial extent is much larger than that of a sensing electrode contact at location *x*_0_ (or, more precisely, that of its *ϕ* along the fiber; see as well Sec. 3.1), *S*(*t*) reduces to:

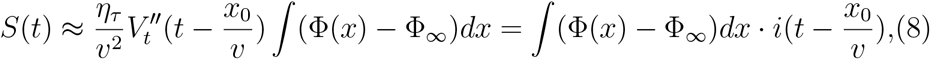

which is proportional to the time-varying transmembrane current at the electrode location (see Appendix A.2 for full derivation). This is a common situation for myelinated fibers.

Similarly (see Appendix A.2), when the AP’s spatial extent is much shorter than that of the sensing electrode contact:

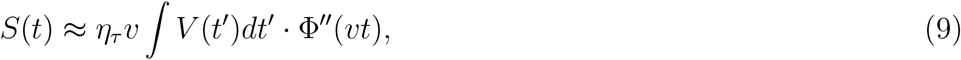

which is proportional to the recording electrode activating function [20] at the propagating location of the AP (i.e., *S*(*t*) does not reveal information about the its shape). In summary, unless the characteristic length scales of the recording electrode potential and the AP are comparable, zeroth and first order contributions vanish (see 4.1), and only small second order contributions remain. This is a consequence of the cancellation of the positive and negative lobes of *V* ^*′′*^, respectively *ϕ*^*′′*^.

#### 4.1.2. Fiber Population

Furthermore, additional signal cancellation occurs on the fiber population level, as the SFAP contributions consist of positive and negative lobes that mostly cancel out when integrated over time (apparent, e.g., when integrating over Eqn.(8) or (9)). The fiber-diameter dependent conduction velocity leads to shifted SFAP arrival times, such that low-order eCAP contributions from fiber populations with smoothly varying diameters cancel. Higher order, diameter-dependent contributions such as degree of recruitment, diameter probability density, and signal contribution scaling are primarily responsible for the eCAP signal. This can make the eCAP signal complex to interpret and sensitive to details of the topological fiber distribution and nerve anatomy.

We have shown that in many fascicles, a significant fraction of fibers, with small diameters, are not recruited (Fig. 5.c-d). In most fascicles, the recruitment curve is steep (Fig. 5.d). Because conduction velocity is proportional to fiber diameter, SFAPs from smaller diameter fibers arrive later than those of larger diameter fibers. When there is a sharp cutoff in the diameter of the recruited fibers, the SFAP contributions of the activated fibers above the cutoff are not compensated anymore. This leads to large-amplitude deflections in the portion of the eCAP associated with the arrival of APs from the fibers immediately above the cutoff (Fig. 5.e), which in turn results in non-monotonicity in the peak-to-peak amplitude of the eCAP as a function of stimulus current (Fig. 6). To our knowledge, the impact of partial fascicular recruitment, have not been previously reported and critically influences the interpretation of the eCAP.

### 4.2. Limitations and Issues

#### 4.2.1. Extended Reciprocity Theorem

The only assumption underlying the derivation of the extended reciprocity theorem is that the totality of the current sources cancel (i.e., charge is conserved). Besides the physical justification for that condition, it is also evident that dropping this condition results in a lack of well-definedness, as the measurable signal would become dependent on the chosen electric potential Gauge (i.e., *ϕ*(*x*) → *ϕ*(*x*) + *ϕ*_0_ changes *S*(*t*)).

By computing the current source density as proportional to the second derivative of the transmembrane potential *V*_*x*_, its integral becomes proportional to the difference between the spatial gradient of *V*_*x*_ at the two terminals. If non-physical boundary conditions are used in the electrophysiology simulations or the multiplying potential *ϕ* is non-zero where the fiber model is truncated, care must be taken to include corresponding sinks to balance the current in the model.

#### 4.2.2. Semi-Analytical eCAP Model

The semi-analytical nerve eCAP model makes several assumptions that may limit its applicability. We assume that the temporal shape of the AP is independent of diameter. While this assumption has been validated for the fiber models used in this study (Fig. B1.c-d), it may not hold *in vivo*. We further assume that there is no systematic or stochastic background nerve activity. The model must also be refined, when complex stimulation paradigms are applied, such as multi-polar pulse-shapes, pulse-trains with high repetition rates, or high stimulation intensities that result in conduction blocking and non-monotonous recruitment curves. Assuming a reciprocal dependence of stimulation thresholds on fiber diameter is not uniformly correct for all fiber types and stimulation exposure lengths [21]. When it fails to hold, recruitment curves must instead be determined for a population of fibers representative of their real distribution. We find that applying a Gaussian jitter to the recruitment threshold for each fiber smooths the eCAP (Fig. G1).

For myelinated fibers, transmembrane current is mostly restricted to Ranvier nodes. When the exposing potential *ϕ* varies significantly over the length-scale of inter-node distances, it could become necessary to account for the spatial discreteness of the transmembrane current sources (incl. their stochastic position variability) and to replace the second derivative in 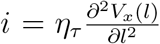 by its finite-difference equivalent, while adapting *η*_*τ*_ to compensate for the reduced current source extent.

The finite element model assumes perfect contact between the recording electrodes and the nerve, and does not account for capacitive effects or scar tissue formation due to the insertion of a foreign body. The generalizability of the finite element model is also limited by uncertainty about the dielectric tissue properties and the perineurium thicknesses.

In principle, a too-coarse sampling of fiber diameters would limit the validity of our results. However, we show that doubling the sampling density, from 2000 to 4000 discrete diameters, hardly affects the signal (Fig. H1).

#### 4.2.3. Nerve Topology Differences Between Simulation And Measurements And Inter/Intra-Subject Variability

In view of the sensitive dependence of the eCAP signal on details of the topological fascicle and fiber distribution, as well as the nerve anatomy, it is important to keep in mind that the porcine histological data was not acquired from the same animal as the eCAP data. Thus, important differences can be expected, complicating validation. In view of future therapeutic application, this raises the problem of how to deal with important inter/intra-subject variability of the vagus topology. The following approaches appear to be relevant: 1) include minimal inter/intra-subject variability in the criteria for implantation site selection, 2) develop non-invasive means of characterizing subject- and implantation-site-specific topologies and fiber distributions (potentially by analyzing eCAP or electrical impedance tomography (EIT) signals; c.f. [22]), or 3) leverage closed-loop control strategies to compensate for the response prediction uncertainty.

#### 4.2.4. Unmyelinated Fiber Contributions

Our model produces a substantially larger signal from unmyelinated fibers than observed experimentally. This is mostly due to the presence of large (*>* 1 *µm*) unmyelinated fibers in the model. Other studies have observed a much smaller number of fibers in this size range [23], albeit in humans rather than pigs. As the fiber diameter distribution in the experiment is unknown, it may be that differences between that distribution and our reference distribution explain differences in the calculated signal.

It is also possible that the observed discrepancies may be due to the choice of unmyelinated fiber model. Models of unmyelinated fibers differ greatly, and have not been extensively validated. While myelinated fiber models are more well-established, the use of a different myelinated fiber model may also have an impact on our results.

### 4.3. Comparison With Previous Work

Our model differs from that of Peña et al. [10] in key aspects: 1) It considers fiber recruitment and accounts for partial activation, which is revealed to be crucial in interpreting eCAP signals. 2) It studies multi-fascicular porcine VNs, rather than the monofascicular rabbit nerve. 3) It is amenable to analytical evaluation, which permits the establishment of the relationship between signal features and parameters of interest. A technical differences is our use of simulated transmembrane voltages vs. transmembrane currents, which avoids separate computations for different fiber diameters. In conclusion, these developments allow for application to complex, multifascicular nerves and neurovascular bundles, and avoid resorting to problematic simplifications, such as neglecting the diameter dependence of partial fascicular activation.

### 4.4. Future Directions

We anticipate that our model can be used to solve the inverse problem of reconstructing the distribution of fiber diameters in a nerve. While the overall distribution of diameters is known, the relative fractions of each fiber type in each fascicle cannot be known for an individual. However, if the location of the fascicles can be reconstructed, for example using EIT, the fiber proportions in each fascicle can be obtained by optimizing the agreement between measured and modeled eCAP signal. Knowledge of the fiber type distribution over fascicles for an individual may be useful for interpreting eCAPs signal or for maximizing stimulation selectivity and targeting, e.g., in treatment planning.

## 5. Conclusions

We have developed an extended reciprocity theorem approach suitable for the efficient computation of signals resulting from spatially extended electrophysiological activity in complex anatomical and dielectric environments. It has been used to establish a simplified analytical SFAP model – revealing interesting differences between myelinated fiber and unmyelinated fiber SFAP behavior – and a semi-analytical eCAP model. Employing these modeling approaches, simulations of VNS and recording were performed that leverage histological information about the multifascicular VN anatomy and its fiber population statistics. The impact on eCAP shape and magnitude of varying stimulation amplitude and stimulus and recording electrode configuration was studied. The eCAP signal is revealed to be more complex than commonly assumed, because of factors, such as cancellation effects on the single fiber and fiber population level and partial fiber recruitment causing strong signal contributions. This complexity on the one hand complicates signal interpretation, but on the other hand might offer the opportunity to maximize signal information content by adapting the eCAP recording setup to increase the sensitivity to the quantity- or features-of-interest. The analytical and semianalytical model are valuable instruments for that purpose, as they help understand the impact of electrode geometry (e.g., contact separation, dimension, orientation, mono-vs. bipolar, location) on eCAP recordings. The ratio of the width of the AP to the extent of the potential associated with the electrode is revealed to be a key parameter.

Our methods can be used to rapidly assess new stimulation and recording setups involving complex nerves and neurovascular bundles, e.g., for closed-loop control in bioelectronic medicine applications.

## 6. Acknowledgements

NeuHeart received funding from the European Union’s Horizon 2020 research and innovation programme under grant agreement No 824071.

This study was supported by funding to the Blue Brain Project, a research center of the École polytechnique fédérale de Lausanne (EPFL), from the Swiss government’s ETH Board of the Swiss Federal Institutes of Technology.

The authors declare no conflicts of interest.

## Appendix A. Derivation of the Mathematical Model

### Appendix A.1. Extended Reciprocity Theorem

We represent a neural fiber as a line with a transmembrane current density *i*(*l*) along its length. For simplicity, we will first consider discretized point current sources *I*_*i*_, which can either represent the integrated current density *i*(*l*_*j*_) · Δ*l* over the *i*-th line segment of length Δ*l*, or the current leaving a Ranvier node separated by a distance Δ*l* from the next unmyelinated location on a myelinated fiber. It can be proven by induction that placing small dipolar sources 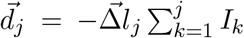 between the locations *x*_*i*_ and *x*_*i*+1_ of two consecutive current sources separated by 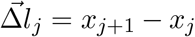 reproduces the series of current sources *I*_*j*_ (charge conservation ensures that the last current source at the terminal is correctly obtained). Applying the reciprocity theorem, we get:

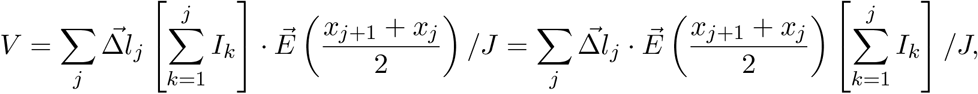

which in the continuum limit becomes:

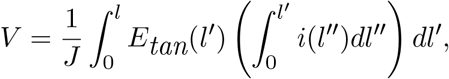

where the tangential field 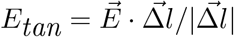 Using integration by parts we obtain:

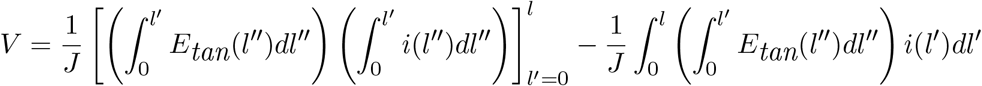

Using the definition of the electric potential, 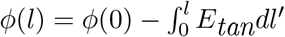

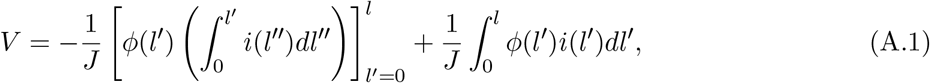

Because of current conservation, 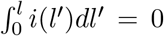 and the expression reduces to the *extended reciprocity theorem*, Eqn.(2):

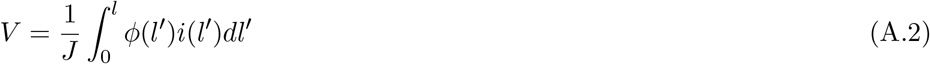

With Φ = *ϕ/J*,

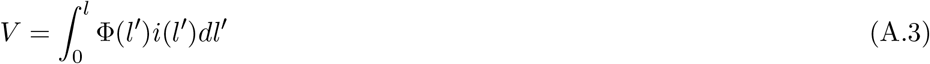

### Appendix A.2. Analytical eCAP Contribution Model

From the extended reciprocity theorem, it follows that the SFAP is the convolution in space of the transmembrane current with the extracellular potential along the neuron:

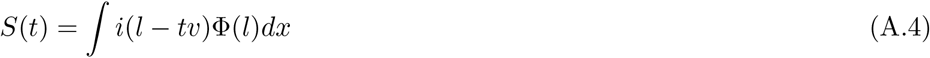

The trans-membrane current can be obtained as 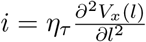, where 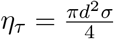, *σ* is the axial conductance of an axon, *V*_*x*_ is the transmembrane voltage (distinguished by the subscript *x* from the temporal transmembrane potential *V*_*t*_(*t*)), and *d* is its diameter [12]. Thus,

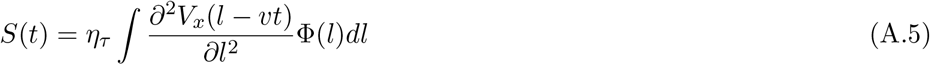

where Φ is the extracellular potential generated by the recording electrodes, normalized to the applied current, and *v* is the conduction velocity.

If *V*_*x*_(*x*) is the transmembrane potential along the fiber, then, for *V*_*x*_ at t = 0, 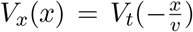, where *V*_*t*_(*t*), obtained at *x* = 0, is the profile of the AP in time that has the benefit of only being weakly fiber diameter dependent. For convenience, we define *t* = 0 to coincide with the peak of *V*_*t*_. Thus, 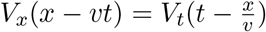, where the important diameter dependence enters through *v*(*d*). With a change of variables *x* = *t*^*′*^ · *v*, we can express this convolution in the time domain (Eqn.(3)):

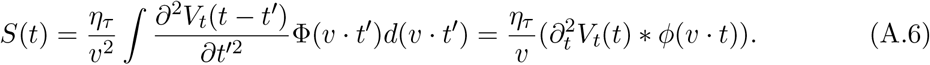

For propagating APs, Eq. A.4 can be expressed as Eq. A.7:

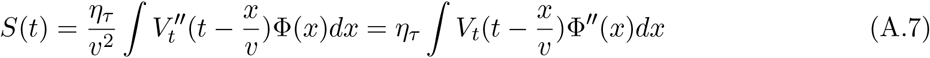

If the AP’s spatial extent is much larger than that of a sensing electrode contact at location *x*_0_ (or, more precisely, that of its Φ along the fiber; see as well Sec. 3.1), 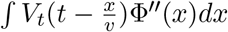 can through Taylor expansion be approximated to second order as:

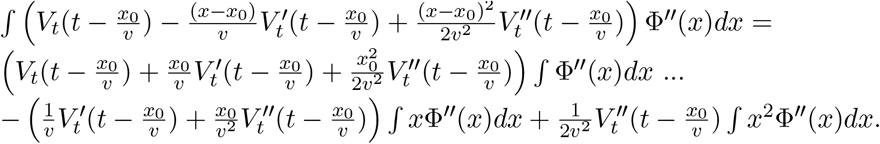

This is a common case for myelinated fibers. By partial integration:

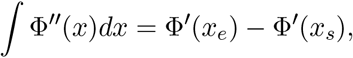

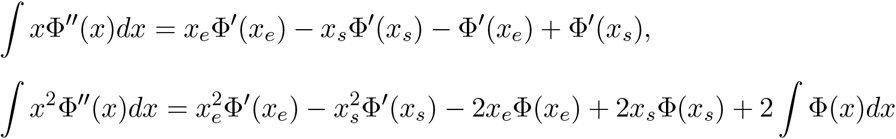

At sufficiently remote terminals *x*_*s*_ and *x*_*e*_, Φ^*′′*^(*x*_*e*_) = Φ^*′*^(*x*_*e*_) = Φ^*′*^(*x*_*s*_) = Φ^*′*^(*x*_*s*_) = 0 and Φ(*x*_*e*_) = Φ(*x*_*s*_) = Φ_∞_, such that *S*(*t*) reduces to:

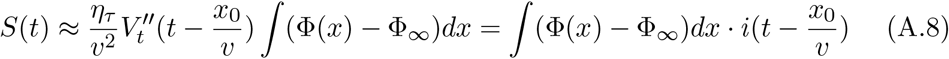

For the converse case, when the spatial extent of the AP is much shorter than that of the electrode potential, a similar analysis shows that 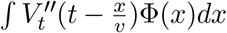 can be approximated as:

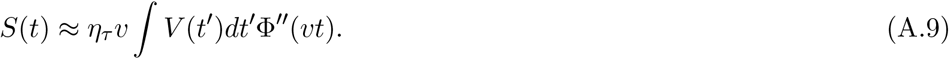

To obtain this result, we substitute *x* = *v* · *t*^*′*^, expand Φ^*′′*^(*x*) around *v* · *t* to second order, and consider that 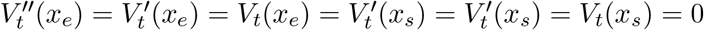.

## Appendix B. Fiber Model Properties

As discussed in Sec. 2.1.4, SFAPs for each of the sampled diameters are rescaled and summed to synthesize the eCAP. Rescaling is appropriate, because the temporal AP shape *V*_*t*_ only weakly depends on *d* (see Fig. B1.c-d). For myelinated fibers, conduction velocity is proportional to fiber diameter *v*_*m*_(*d*) = *a* · *d*, whi<u>le</u> for unmyelinated fibers, it is proportional to the square root of fiber diameter 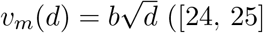 [24, 25]; see Fig. B1.a-b).

## Appendix C. *In Vivo* Experiments

The protocol for all animal studies (no. 76/2014 PR) was approved by the Italian Ministry of Health and complied with Italian law (D.lgs. 26/2014). This study involved five healthy adult male Göttingen minipigs (Ellegaard Göttingen Minipigs A/S, Dalmose, Denmark; average body weight of 35 kg). The animals were premedicated with Zoletil® (10 mg/kg) and Stressnil (1 mg/kg). Anesthesia was induced with propofol (2 mg/kg intravenously) and maintained with 1% to 2% sevoflurane in air enriched with 50% oxygen during mechanical ventilation [26][27]. To prevent dehydration, each animal received a continuous infusion of 500 mL of sodium chloride (0.9%) solution throughout the experiments. A longitudinal incision followed by a sternotomy was performed, and blunt dissection was used to expose the VN at both the cervical and thoracic levels. Differential recordings from the four channels commercial cuff electrode (World Precision Instruments, Sarasota, FL, USA) were acquired using a TDT system (Tucker Davis Technologies; Alachua, FL, USA), with a sampling rate of 24k samples/s. Each recording session was composed of 10 s baseline activity, followed by 1 min VNS, and 1 min of washout.

## Appendix D. EM Modeling

The finite element simulation solves the equation ∇*σ*∇*ϕ* = 0, where *σ* is the electrical conductivity distribution and Φ is the electric potential, from which the E-field is obtained as 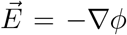 and the current density as 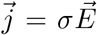. This approximation is suitable, as ohmic currents dominate over displacement currents and the simulation domain is much smaller than the wavelength in tissue [28]. Material properties were assigned to all tissues and materials according to Table D1, including the anisotropy of the intrafascicular endoneurium (tensorial *σ*). Thin resistive layer settings were assigned to patches at the interfaces between fascicles and the interfascicular tissue with a thickness computed as 3 % of the fascicle’s equivalent diameter [29].

**Figure B1.**
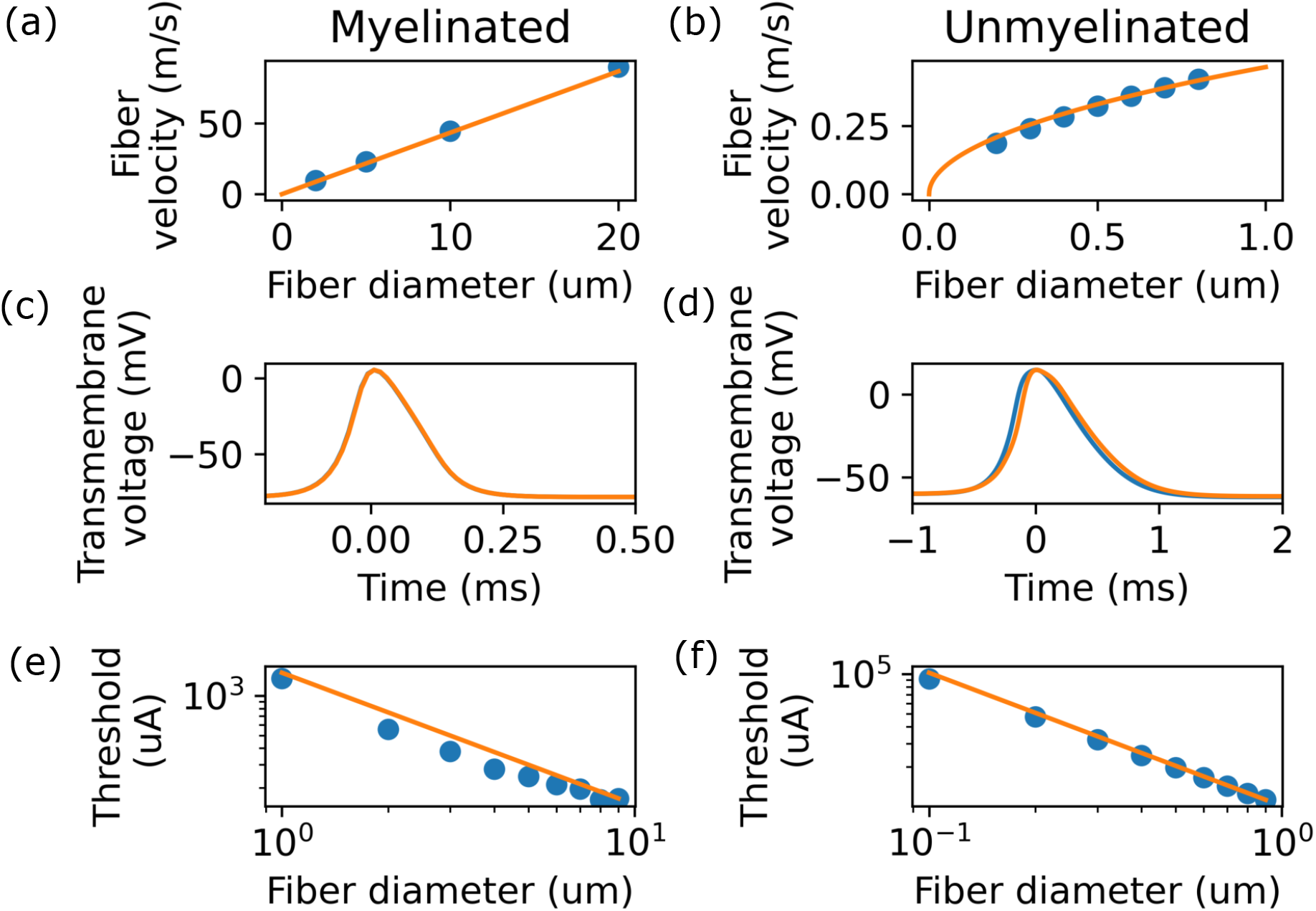
First row: Fiber velocity is linear with respect to diameter for myelinated fibers (panel a) and proportional to the square root of velocity for unmyelinated fibers (panel b). Blue dots indicate velocities determined by simulation; orange lines indicate expected relationships. Second row: Temporal shape of the AP does not vary significantly with diameter for myelinated fibers (panel c; orange: *d* = 10 *µ*m, blue: *d* = 20 *µ*m) and for unmyelinated fibers (panel d; orange: *d* = 0.4 *µ*m, blue: *d* = 0.8 *µ*m). Third row: Stimulus threshold is inversely proportional to fiber diameter for myelinated (panel e) and unmyelinated (panel f) fibers. Blue dots indicate thresholds determined by simulations; orange lines indicate expected relationship.

As described in Section 2.1.4, the sensitivity function Φ_*i*_(*l*) is calculated for each fascicle, by interpolating the potential field obtained from the finite element simulation along the fascicle center-line. The interpolated potential field is post-processed to ensure that it reaches 0 at the ends of the nerve. At each end of the Φ_*i*_(*l*) curve, we identify the point having a predefined minimal slope. We linearize the Φ curve from this point, until the intersection with the line Φ = 0, after which Φ is set to 0.

To simulate the electrode placement along the major axis of the nerve, the insulating substrate is excluded, in order to prevent the electrode from piercing any of the fascicles. As all of the contacts, on both sides of the electrode, are active in our simulation, the exclusion of the substrate does not have a signficant effect on the resulting E-field.

**Table D1.**
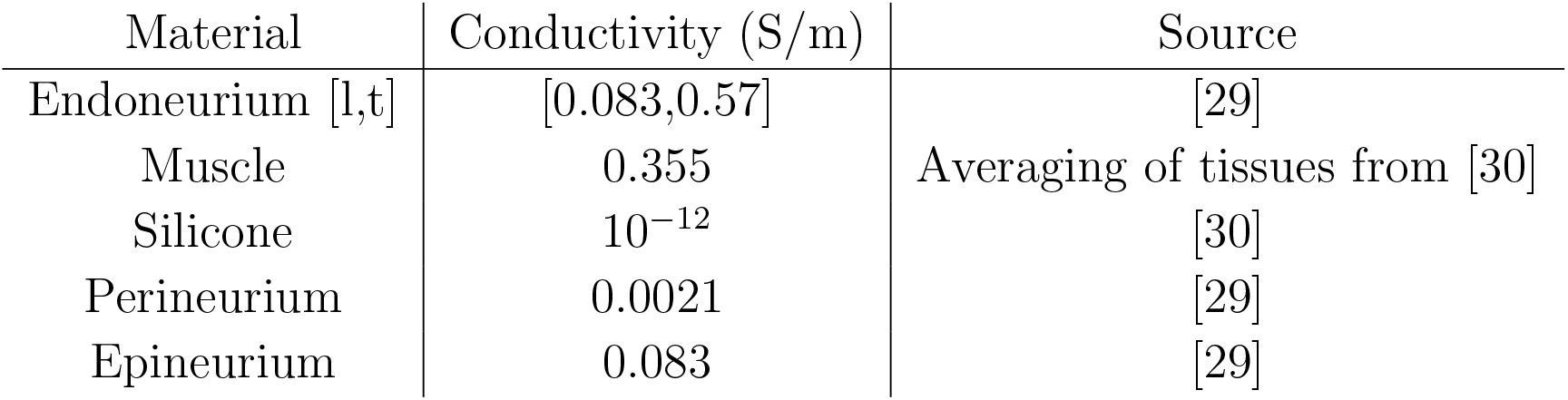
Tissue conductivities. For anisotropic endoneurium, the longitudinal and transversal conductivities are listed.

## Appendix E. Neuronal Dynamics Simulations

Fiber electrophysiology simulations were executed within Sim4Life v8.0 using the T-Neuro module, which couples the EM solver with Yale’s NEURON [31]. Reference simulations to obtain shapes of the AP were performed with an axon diameter of 20 *µ*m for myelinated fibers and 0.8 *µ*m for unmyelinated fibers, using a time-step of 0.0025 ms. Simulations performed with myelinated and unmyelinated diameters of 10 *µ*m and 0.4 *µ*m, respectively,confirmed that the shape of the AP is independent of fiber diameter (Fig. B1.c-d). For neural titration simulations, *d*_0_ = 0.8 *µm* for unmyelinated fibers and 4 *µm* for myelinated fibers.

## Appendix F. Implementation

The semi-analytic model was implemented on the Blue Brain Project’s BB5 supercomputer, housed at the Swiss National Supercomputing Center datacenter. Simulations were run using 39 nodes, with two 2.30 GHz, 18 core Xeon SkyLake 6140 CPUs, and 382 GB DRAM each. Simulations were completed in under 5 min. Finite element models were run on a Windows server with a 2.30 GHz, 32-core Xeon E5-4610 CPU and 512 GB RAM. Only one core was used. Neural simulations were run on a similar server, using 55 cores.

The finite element models used in this paper are accessible at https://osparc.io/#/study/e03935d2-79a5-11ef-8269-0242ac174ae0. The code used to run the semi-analytic model and make the figures are available at https://github.com/joseph-tharayil/vagusNerve.

## Appendix G. Gaussian Jitter in Recruitment Curves

For each fiber in the neural titration simulation, we add a normally-distributed random variable to the threshold current, with mean of 0 and standard deviation equal to a particular fraction *f* of the threshold current. Then, as described in Section 2.3.3, the recruitment probability for each fiber diameter in the semi-analytic model is parameterized based on the (now adjusted) threshold values from the titration. The process is repeated for *f* ∈ *{*0.1, 0.2, 0.3, 0.4*}*

By applying Gaussian jitter to the recruitment threshold for each fiber, the eCAP signal is smoothed(Fig. G1).

**Figure G1.**
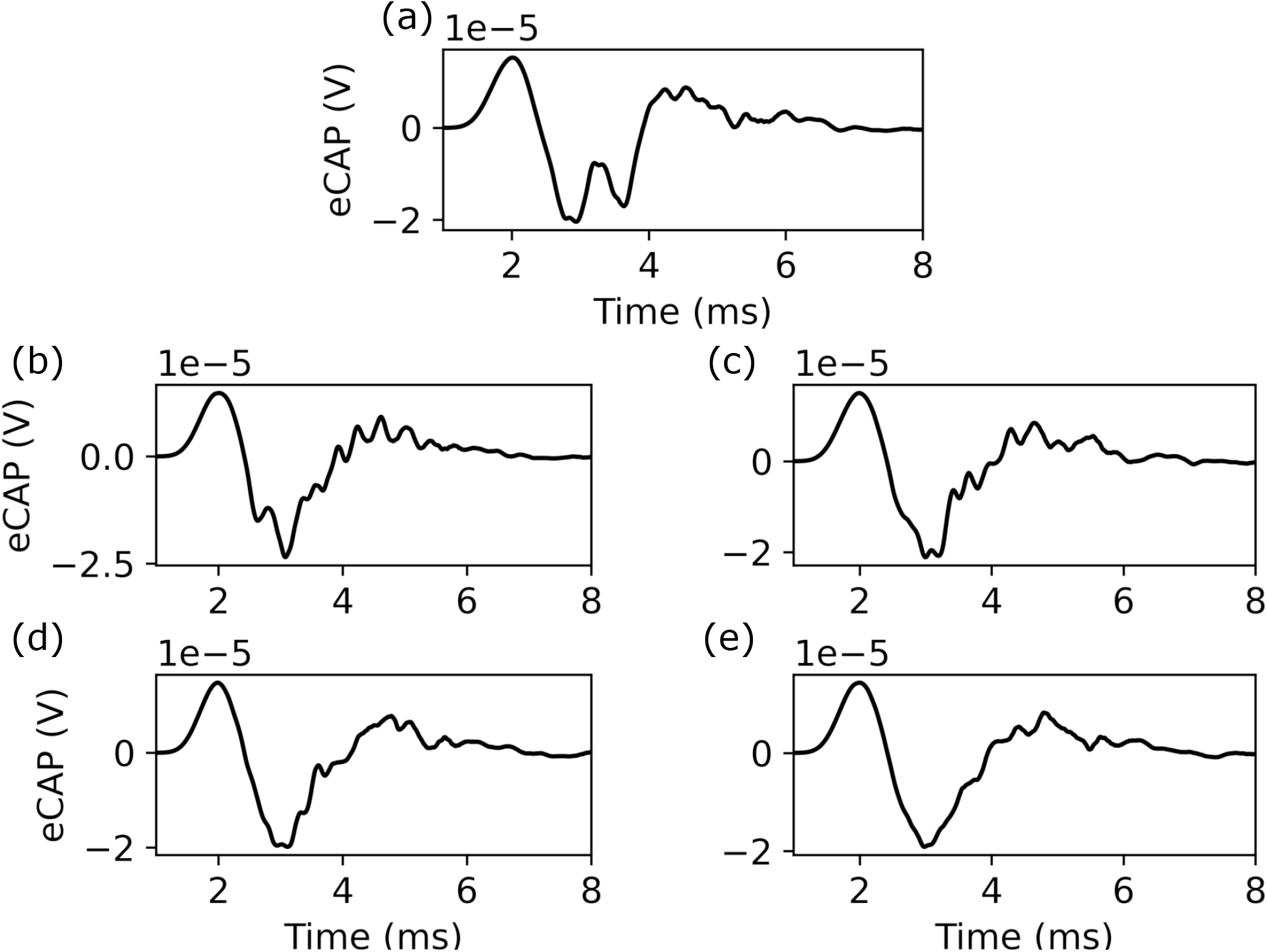
a: eCAP produced at 31.25 *µ*A without adding Gaussian jitter to the recruitment curve. b-e: eCAPs produced at 31.25 *µ*A stimulus current with Guassian jitter of 10%, 20%, 30%, and 40%, respectively, added to the recruitment curve.

## Appendix H. No Effect of Diameter Sampling Density on eCAP

To ascertain whether we have sufficient diameter sampling resolution in our implementation of Equation 6, we simulated eCAPs with 2000 and 4000 diameters sampled between 0.1 and 15 *µ*m. The eCAPs are almost identical.

## Appendix I. Recording Configuration Critically Influences Signals

Compared to monopolar recordings, biopolar ones with small recording contact separation correspond to a finite difference derivative, such that the number of signal phases with alternating polarity (‘lobes’) is increased by 1 – for the simulated eCAP in Fig. I1.a the monopolar signal is approximately biphasic and the bipolar one triphasic. As the contact separation increases, this corresponds to coarsening and smoothing of the finite difference derivative. It results in reduced fluctuations and noise (Fig. I1.b), which potentially represents a loss in temporal information content. If the separation is increased further, the signal shape undergoes a complex transformation, until two distinct monopolar recording shapes emerge.

**Figure H1.**
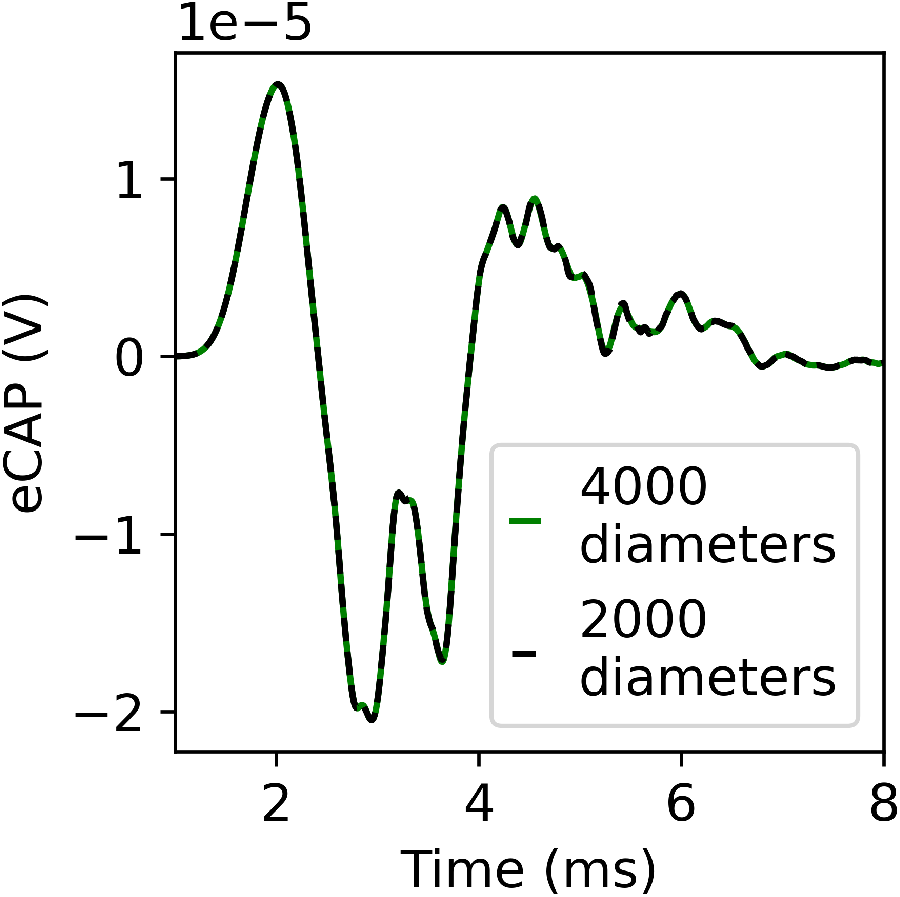
Increasing the diameter resolution in the discretized eCAP computation from 2000 to 4000 hardly affects the modeling results.

**Figure I1.**
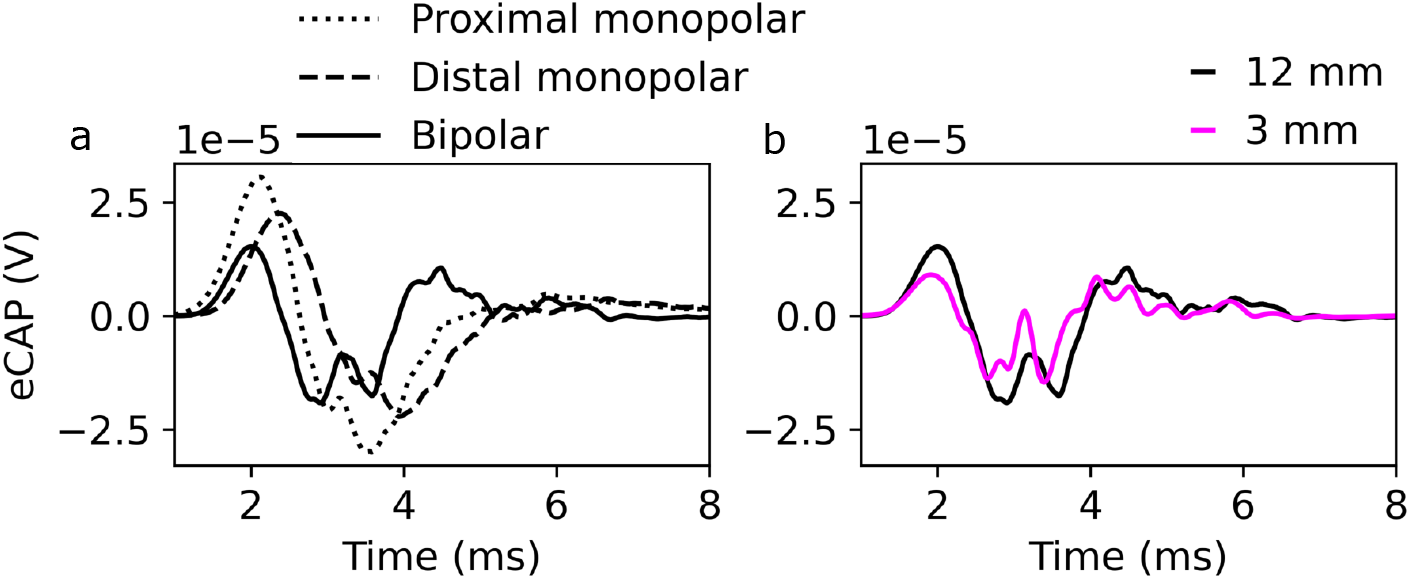
The recording configuration affects eCAPs recorded by a four contact cuff electrode (6 cm from the stimulus site, 31.25 *µ*A stimulus current). a: Monopolar recordings with the most proximal or the most distal contact (distant reference), plotted alongside a bipolar recording using both contacts (12 mm separation). b: Bipolar recordings using the outermost or innermost electrodes (12 vs. 3 mm separation).

## References

[1] R. S. Wijesinghe, F. L. H. Gielen, and J. P. Wikswo, “A model for compound action potentials and currents in a nerve bundle i: The forward calculation,” Annals of Biomedical Engineering, vol. 19, no. 1, pp. 43–72, 1991. [Online]. Available: http://link.springer.com/10.1007/BF02368460

[2] I. Gupta, A. M. Cassara, I. Tarotin, M. Donega, J. A. Miranda, D. M. Sokal, S. Ouchouche, W. Dopson, P. Matteucci, E. Neufeld, M. A. Schiefer, A. Rowles, P. McGill, J. Perkins, N. Dolezalova, K. Saeb-Parsy, N. Kuster, R. F. Yazicioglu, J. Witherington, and D. J. Chew, “Quantification of clinically applicable stimulation parameters for precision near-organ neuromodulation of human splenic nerves,” Communications Biology, vol. 3, no. 1, p. 577, 2020.

[3] H. Yuan and S. D. Silberstein, “Vagus nerve and vagus nerve stimulation, a comprehensive review: Part ii,” Headache: The Journal of Head and Face Pain, vol. 56, no. 2, pp. 259–266, 2016. [Online]. Available: https://headachejournal.onlinelibrary.wiley.com/doi/abs/10.1111/head.12650

[4] M. J. Capilupi, S. M. Kerath, and L. B. Becker, “Vagus Nerve Stimulation and the Cardiovascular System,” Cold Spring Harb Perspect Med, vol. 10, no. 2, Feb 2020.

[5] F. Agnesi, C. Zinno, I. Strauss, A. Dushpanova, V. Casieri, F. Bernini, D. Terlizzi, K. Gabisonia, V. Paggi, S. P. Lacour, V. Lionetti, and S. Micera, “Cardiovascular response to intraneural right vagus nerve stimulation in adult minipig,” Neuromodulation: Technology at the Neural Interface, 2023. [Online]. Available: https://www.sciencedirect.com/science/article/pii/S1094715923001319

[6] C. Zinno, F. Agnesi, G. D’Alesio, A. Dushpanova, L. Brogi, D. Camboni, F. Bernini, D. Terlizzi, V. Casieri, K. Gabisonia, L. Alibrandi, C. Grigoratos, J. Magomajew, G. D. Aquaro, S. Schmitt, P. Detemple, C. M. Oddo, V. Lionetti, and S. Micera, “Implementation of an epicardial implantable MEMS sensor for continuous and real-time postoperative assessment of left ventricular activity in adult minipigs over a short- and long-term period,” APL Bioengineering, vol. 8, no. 2, p. 026102, 04 2024. [Online]. Available: 10.1063/5.0169207

[7] M. Haberbusch, S. Frullini, and F. Moscato, “A numerical model of the acute cardiac effects provoked by cervical vagus nerve stimulation,” IEEE Transactions on Biomedical Engineering, vol. 69, no. 2, pp. 613–623, 2021.

[8] M. Haberbusch, B. Kronsteiner, A.-M. Kramer, A. Kiss, B. K. Podesser, and F. Moscato, “Closed-loop vagus nerve stimulation for heart rate control evaluated in the langendorff-perfused rabbit heart,” Scientific Reports, vol. 12, no. 1, p. 18794, 2022.

[9] M. Haberbusch and E. Neufeld, “From models to heartbeats: Computational design of vagus nerve stimulation for cardiac health,” Webinar, October 2023. [Online]. Available: https://www.physiology.org/detail/event/2023/10/18/default-calendar/webinar-series-from-models-to-heartbeats-computational-design-of-vagus-nerve-stimulation-for-cardiac-health?

[10] E. Peña, N. A. Pelot, and W. M. Grill, “Computational models of compound nerve action potentials: Efficient filter-based methods to quantify effects of tissue conductivities, conduction distance, and nerve fiber parameters,” PLOS Computational Biology, vol. 20, no. 3, pp. 1–35, 03 2024. [Online]. Available: 10.1371/journal.pcbi.1011833

[11] J. Tharayil, S. Farcito, B. Lloyd, T. Newton, C. Zinno, F. Agnesi, A. Cassara, E. Neufeld, S. Micera, and N. Kuster, “A computational model of porcine vagus nerve stimulation and compound action potential sensing for enhanced closed-loop cardiac control,” North American Neuromodulation Society Annual Conference, January 2024.

[12] M. S. Spach, R. C. Barr, G. A. Serwer, J. M. Kootsey, and E. A. Johnson, “Extracellular potentials related to intracellular action potentials in the dog purkinje system,” Circulation Research, vol. 30, no. 5, pp. 505–519, 1972. [Online]. Available: https://www.ahajournals.org/doi/abs/10.1161/01.RES.30.5.505

[13] M. Ikeda and Y. Oka, “The relationship between nerve conduction velocity and fiber morphology during peripheral nerve regeneration,” Brain Behavior, vol. 2, no. 4, pp. 382–390, 2012.

[14] G. Matsumoto and I. Tasaki, “A study of conduction velocity in nonmyelinated nerve fibers,” Biophysical Journal, vol. 20, no. 1, pp. 1–13, 1977. [Online]. Available: https://www.sciencedirect.com/science/article/pii/S000634957785532X

[15] N. Jayaprakash, W. Song, V. Toth, A. Vardhan, T. Levy, J. Tomaio, K. Qanud, I. Mughrabi, Y.-C. Chang, M. Rob, A. Daytz, A. Abbas, Z. Nassrallah, B. T. Volpe, K. J. Tracey, Y. Al-Abed, T. Datta-Chaudhuri, L. Miller, M. F. Barbe, S. C. Lee, T. P. Zanos, and S. Zanos, “Organ- and function-specific anatomical organization of vagal fibers supports fascicular vagus nerve stimulation,” Brain Stimulation, vol. 16, no. 2, pp. 484–506, 2023. [Online]. Available: https://www.sciencedirect.com/science/article/pii/S1935861X23016856

[16] I. Strauss, F. Agnesi, C. Zinno, A. Giannotti, A. Dushpanova, V. Casieri, D. Terlizzi, F. Bernini, K. Gabisonia, Y. Wu, D. Jiang, V. Paggi, S. Lacour, F. Recchia, A. Demosthenous, V. Lionetti, and S. Micera, “Neural stimulation hardware for the selective intrafascicular modulation of the vagus nerve,” IEEE Transactions on Neural Systems and Rehabilitation Engineering, vol. 31, pp. 4449–4458, 2023.

[17] S. Raspopovic, M. Capogrosso, and S. Micera, “A computational model for the stimulation of rat sciatic nerve using a transverse intrafascicular multichannel electrode,” IEEE Transactions on Neural Systems and Rehabilitation Engineering, vol. 19, no. 4, pp. 333–344, 2011.

[18] J. R. Schwarz and G. Eikhof, “Na currents and action potentials in rat myelinated nerve fibres at 20 and 37 ?c,” Pflügers Archiv, vol. 409, no. 6, pp. 569–577, Aug. 1987. [Online]. Available: 10.1007/BF00584655

[19] D. Sundt, N. Gamper, and D. B. Jaffe, “Spike propagation through the dorsal root ganglia in an unmyelinated sensory neuron: a modeling study,” Journal of Neurophysiology, vol. 114, no. 6, pp. 3140–3153, 2015, pMID: 26334005. [Online]. Available: 10.1152/jn.00226.2015

[20] F. Rattay, “Analysis of models for external stimulation of axons,” IEEE transactions on biomedical engineering, vol. 10, pp. 974–977, 1986.

[21] T. Newton, J. Ordonez, E. Neufeld, and N. Kuster, “Optimizing spinal cord stimulation using a novel green’s function-based generalized activating function,” Neuromodulation, vol. 26, no. 4, p. S44, 2023.

[22] C. A. Chapman, K. Aristovich, M. Donega, C. T. Fjordbakk, T.-R. Stathopoulou, J. Viscasillas, J. Avery, J. D. Perkins, and D. Holder, “Electrode fabrication and interface optimization for imaging of evoked peripheral nervous system activity with electrical impedance tomography (eit),” Journal of Neural Engineering, vol. 16, no. 1, p. 016001, 2018.

[23] L. A. Havton, N. P. Biscola, E. Stern, P. V. Mihaylov, C. A. Kubal, J. M. Wo, A. Gupta, E. Baronowsky, M. P. Ward, D. M. Jaffey, and T. L. Powley, “Human organ donor-derived vagus nerve biopsies allow for well-preserved ultrastructure and high-resolution mapping of myelinated and unmyelinated fibers,” Sci Rep, vol. 11, no. 1, p. 23831, Dec 2021.

[24] G. Matsumoto and I. Tasaki, “A study of conduction velocity in nonmyelinated nerve fibers,” Journal of Biophysics, vol. 20, no. 1, pp. 1–13, Oct. 1977.

[25] S. Waxman and M. Bennett, “Relative conduction velocities of small myelinated and nonmyelinated fibres in the central nervous system,” Nature New Biology, vol. 238, no. 85, pp. 217–219, Aug. 1972.

[26] D. Ferraro, G. D’Alesio, D. Camboni, C. Zinno, L. Costi, M. Haberbusch, P. Aigner, M. Maw, T. Schlöglhofer, E. Unger, A. Aliperta, F. Bernini, V. Casieri, D. Terlizzi, G. Giudetti, J. Carpaneto, G. Pedrizzetti, S. Micera, V. Lionetti, F. Moscato, L. Massari, and C. M. Oddo, “Implantable fiber bragg grating sensor for continuous heart activity monitoring: Ex-vivo and in-vivo validation,” IEEE Sensors Journal, vol. 21, no. 13, pp. 14 051–14 059, 2021.

[27] V. Lionetti, S. L. Romano, G. Bianchi, F. Bernini, A. Dushpanova, G. Mascia, M. Nesti, F. Di Gregorio, A. Barbetta, and L. Padeletti, “Impact of acute changes of left ventricular contractility on the transvalvular impedance: Validation study by pressure-volume loop analysis in healthy pigs,” PLOS ONE, vol. 8, no. 11, pp. 1–9, 11 2013. [Online]. Available: 10.1371/journal.pone.0080591

[28] C. A. Bossetti, M. J. Birdno, and W. M. Grill, “Analysis of the quasi-static approximation for calculating potentials generated by neural stimulation,” Journal of Neural Engineering, vol. 5, no. 1, pp. 44–53, dec 2007.

[29] Y. Grinberg, M. A. Schiefer, D. J. Tyler, and K. J. Gustafson, “Fascicular perineurium thickness, size, and position affect model predictions of neural excitation,” IEEE Transactions on Neural Systems and Rehabilitation Engineering, vol. 16, no. 6, pp. 572–581, Dec 2008.

[30] P. Hasgall, F. Di Gennaro, C. Baumgartner, E. Neufeld, B. Lloyd, M. Gosselin, D. Payne, Klingenböck, and N. Kuster, “IT’IS database for thermal and electromagnetic parameters of biological tissues.”

[31] M. L. Hines and N. T. Carnevale, “Neuron: A tool for neuroscientists,” The Neuroscientist, vol. 7, no. 2, pp. 123–135, 2001, pMID: 11496923. [Online]. Available: 10.1177/107385840100700207

